# Joint analysis of RNA-DNA and DNA-DNA interactomes reveals their strong association

**DOI:** 10.1101/2024.11.30.626180

**Authors:** Dmitry S. Zvezdin, Artyom A. Tyukaev, Anastasia A. Zharikova, Andrey A. Mironov

## Abstract

At the moment, many non-coding RNAs that perform a variety of functions in the regulation of chromatin processes are known. An increasing number of protocols allow researchers to study RNA-DNA interactions and shed light on new aspects of the RNA-chromatin interactome. The Hi-C protocol, which enables the study of chromatin’s three-dimensional organization, has already led to numerous discoveries in the field of genome 3D organization. We conducted a comprehensive joint analysis of the RNA-DNA interactome and chromatin structure across different human and mouse cell lines. We have shown that these two phenomena are closely related in many respects, with the nature of this relationship being both tissue-specific and conserved across human and mouse.

## 1. Introduction

Most of the human genome is transcribed, while the proportion of proteins coding genes is small among these transcripts [1]. The vast majority of transcribed loci are represented by various classes of non-coding RNAs that perform a variety of functions in the nuclei of eukaryotic cells. Small nuclear RNAs are key factors in splicing, small nucleolar RNAs are involved in the processing of ribosomal RNAs, and long non-coding RNAs play diverse roles in chromatin regulation [2]. MALAT1 participates in the organization of nuclear speckles [3], NEAT1 is a key structural element of paraspeckles [4], XIST participates in dose compensation of the second X chromosome of female mammals [5], and FIRRE plays a role in mediating interchromosomal contacts through proteins, thus helping to maintain chromatin structure [6]. In addition, several proteins that play a key role in maintaining the spatial structure of chromatin, such as CTCF and components of the PRC2 protein complex, have the ability to bind RNA, which indicates the possibility of involving RNA molecules in the regulatory function of these proteins [2].

In recent years, a number of methods have been developed to map the RNA-chromatin interactions. First, the methods from the “one-to-all” group appeared (RAP [7], CHIRP [8], CHART [9]), which allow us to study the interactions of one selected RNA with whole genome with good coverage. To be able to map the interactions of all RNAs genome-wide, methods from the “all-to-all” group were developed (Red-C [10], GRID-seq [11], RADICL-seq [12], MARGI [13], iMARGI [14]). In addition to the obvious advantages, they also have a number of disadvantages, namely a high level of noise and very significant incompleteness of data. Despite these limitations, “all-to-all” methods have provided new insights into the RNA-DNA interactome. For example, in the work using the Red-C method, the authors annotated a variety of previously unannotated chromatin associated RNAs (ucaRNA, X RNAs) which had not been detected before, likely due to their strong tendency to localize on chromatin. [10]. Data of the RNA chromatin interactome have a number of biases, in particular, a decrease in the density of contacts with an increase in the distance from the gene encoding the RNA. This shift in the data is caused by contacts of the transcribed RNA (polymerase trace) and the diffusion of RNA through the nucleus. In the future, by analogy with the Hi-C data, we will call this effect - RD-scaling.

Hi-C experimental protocol produces genome-wide mapping of DNA-DNA interactions [15]. Using these data, many discoveries have been made in recent years in the study of the structural organization of chromatin. Various spatial structures formed by chromosomes were annotated on Hi-C maps, such as topologically associated domains (TADs) [16], chromatin loops [17] and A/B compartments [15]. Several studies have examined the relationship between RNA-DNA interactions and chromatin structures, including TADs and A/B compartments. It was found that contacts of most RNAs tend to be localized within the TAD of their gene [11,12]. A preference for interacting with loci in the compartment containing the RNA gene was also observed [12]. However, these analyses were mostly general, and thorough study at the level of individual RNAs was not performed. A large-scale study of the relationship of RNA-DNA interactions and chromatin structure for K562 cells was conducted in 2023 [18]. The barrier effect of the TAD boundaries was experimentally confirmed, and it was also shown that inhibition of transcription and treatment with RNase lead to the formation of new chromatin loops.

In the 2024 study [19], it was shown using the data of the iMARGI protocol that RNA-DNA interactions improve the quality of prediction of Hi-C maps based on DNA sequence with machine learning models. The HiChIRP method was also developed, combining the study of the RNA-DNA interactome and the spatial structure of chromatin [20]. It allows researchers to study DNA-DNA interactions associated with a specific RNA, but it involves mapping of only a limited set of predefined RNAs. Attempts to perform a genome-wide characterization of RNAs involved in regulatory processes in chromatin based on a joint analysis of the RNA-DNA interactome and chromatin structure have not yet been carried out.

In this paper, we propose an approach to the joint analysis of the RNA-DNA interactome and chromatin structure (Figure 1). We show using a number of cell lines that the distribution of RNA contacts across the genome is related to its spatial organization at the intrachromosomal and interchromosomal levels. We demonstrated functional differences in RNA-associated chromatin structures for different RNAs and highlighted the conservation of the RNA-DNA interactome and chromatin structure association across experiments. We also conducted a comprehensive analysis of the relationship between RNA-DNA interaction patterns and certain chromatin structures. Our study shows that many RNAs tend to interact with chromatin loops. We demonstrated that RNAs preferentially interact with the boundaries of their TADs, and the tendency of RNAs to localize outside their parental TAD is partially conserved across human and mouse and dependent on the RNA biotype. The entire analysis was performed using BaRDIC peak caller for RNA-DNA interactions, taking into account the effects of background and RD-scaling [21]. In this study, we adopt the terminology commonly used in studies of chromatin structure: we define cis-contacts as interactions with the chromosome on which the RNA gene is located, and trans-contacts as interactions with other chromosomes.

**Figure 1.**
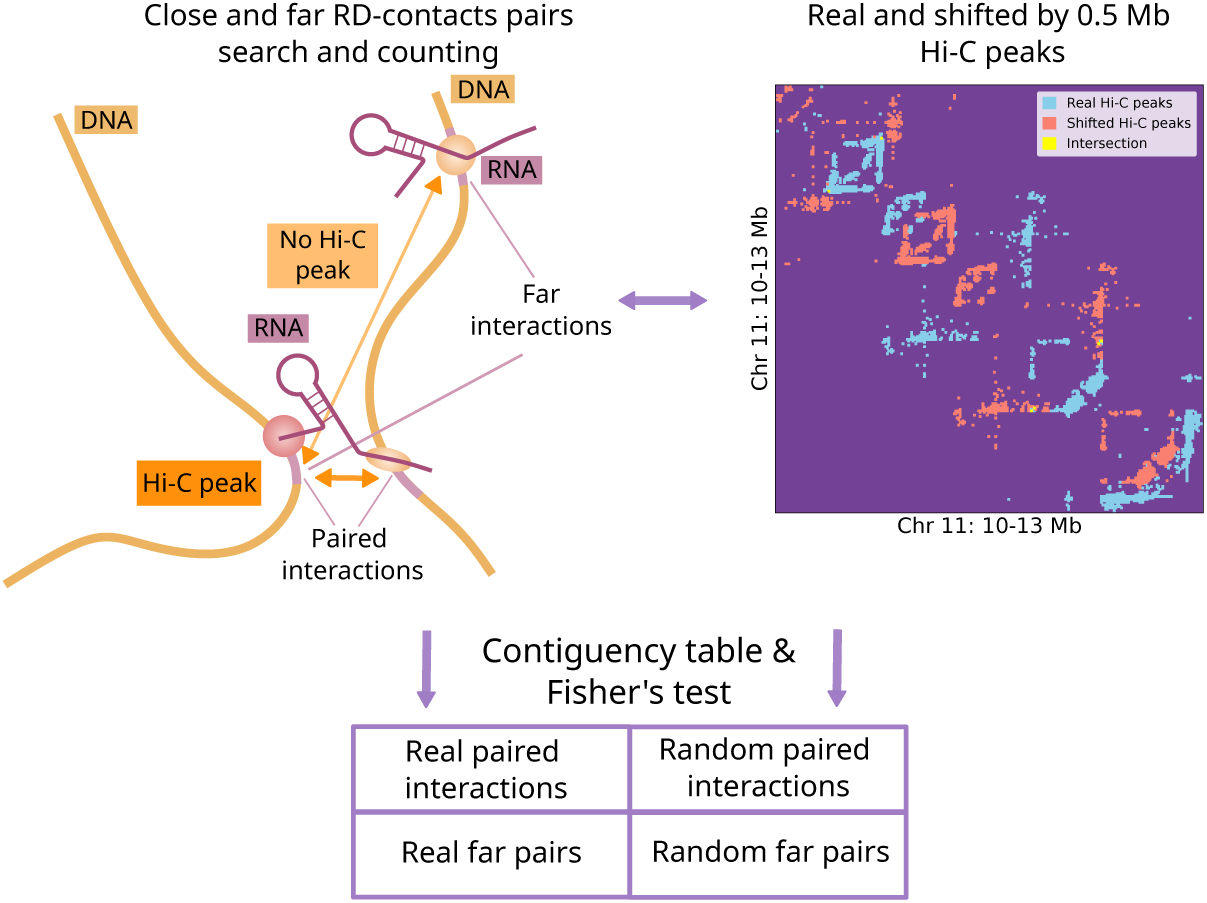
The analysis scheme of the association of chromatin structure and contacts of individual RNAs.

## 2. Materials and Methods

### 2.1. Procedure for identifying associations between RNA-DNA and DNA-DNA interactions

The contact tracks of each individual RNA were binned according to the Hi-C map bin size (10Kb for intrachromosomal interactions and 50Kb for interchromosomal interactions). A pair of RNA-chromatin contacts was considered structurally associated if there was a significant interaction between the corresponding bins of the Hi-C map (based on FitHiC2, Figure 1). To assess whether there was an overrepresentation of such pairs, we used a random model: a Hi-C map shifted along each axis (along the main diagonal for the intrachromosomal case) by a given number of nucleotides, taking into account the A/B compartments (Figure 1, details in Supplementary note; Supplementary Figure S1, Figure S2). In this study, a 2 Mb shift was chosen. Next, the number of such pairs was calculated for each chromosome (for each pair of chromosomes in the case of interchromosomal interactions) and for each RNA independently. The obtained p-values from Fisher’s exact test were adjusted for multiple testing using Benjamini–Hochberg correction (separately for the intra- chromosomal and interchromosomal cases). We considered RNAs with a q-value less than 0.05 to be significantly associated with chromatin structures on a given chromosome (or on a given pair of chromosomes in the case of interchromosomal interaction analysis). Code used for analysis can be found at: https://github.com/ZvezdinDmitry/RNA_DNA_HiC

### 2.2. Data sources

The human gene annotation was compiled from GENCODE (release 35, grch38.p13) [22], small and transport from UCSC [23] and vlinc RNA from the original work on their annotation [24]. For the mouse: GENCODE mm10.p6 and small and transport from UCSC. The annotation of the hypothetical ucaRNAs was taken from the RNA- ChromDB [25] database. Hi-C maps with pre-calculated normalization weights in cool format, as well as the annotation of A/B compartments were taken from the 4DNucleome database (K562: 4DNESI7DEJTM, hESC: 4DNES2M5JIGV, mESC: 4DNESDXUWBD9, mOPC: 4DNESJ9SIAV5). Hi-C data for cell lines with deletions in CTCF RNA-binding domains were taken from the study of the effect of CTCF RNA-binding domain on chromatin structure [26]. Chromatin states ChromHMM annotation was taken from the original work on ChromHMM [27], SPIN states annotation was taken from the work on SPIN states [28].

### 2.3. Data preprocessing

All data manipulations were performed using Python 3.10.13. For intersections and other manipulations with genomic intervals bedtools (v2.31.0) [29] and bioframe (v0.5.0) [30] were used. Only those RNA-DNA contacts that fall into the peaks of the corresponding RNAs were taken for analysis. The peaks were obtained using the program BaRDIC [21] with default parameters for “all-to-all” protocols. The threshold for peak selection was set so that a given proportion of RNA chromatin contacts (10%) fell into peaks (details in Supplementary note). For the analysis of TADs, only intrachromosomal RNA-chromatin contacts were used, so the threshold was selected separately using only them. For “one-to-all” protocols we used the parameters –trans_min 1000 –cis_start 100 –trans_step 100. The threshold was selected similarly to the “all-to-all” data. Contacts within the BaRDIC peaks were used for the analysis. For basic manipulations with Hi-C data, the libraries cooler (v0.9.3) [31] and cooltools (v0.5.4) [32] were used. Significant interactions in the Hi-C data were obtained using the FitHiC2 [33] program for intrachromosomal interactions with a bin size of 10Kb (run separately for each chromosome, FDR: 0.05), and for interchromosomal interactions - 50Kb (FDR: 0.2). Normalization weights were calculated for the maps using the KR algorithm, using the script included in the FitHiC2 package.

### 2.4. Analysis of functional chromatin annotations

The ChromHMM annotation for K562 and hESC cells was converted from the hg18 version of the human genome to the hg38 genome assembly using the liftover [34] web service. ChromHMM and SPIN states were grouped according to tables (Supplementary Table S1, Table S2) to increase their genome coverage and reduce noise level. We considered the SPIN state of Interior_Repr2 separately as a NAD_like, since the authors indicated that these regions tend to aggregate with the nucleolus [28].

To synchronize chromatin annotations and bin pairs of interactions, we used an approach inspired by the work on Multiplex GAM [35]. We assumed that a given bin belongs to a given state if at least one base pair falls within it; therefore, a bin can belong to several different states within the same annotation. A pair of bins is assigned to a given state if both bins are in that state. For each RNA and each chromosome, we independently counted the number of paired contacts between each pair of chromatin states with the same label. To account for the varying proportions of significant Hi-C interactions between pairs of loci, we calculated the expected number of paired contacts for a given chromatin state based on the proportion of Hi-C peaks between loci within that state. The significance of the differences between the observed and expected fractions was assessed using the chi-square test.

### 2.5. Comparative analysis of protocols

To assess how the spatial association of a given RNA is conservative between different experiments, we used Fisher’s exact test with a two-sided alternative hypothesis. We designate one experiment a target experiment, and the other as the reference experiment. The first category in the contingency table corresponds to whether the contacts in the target experiment are paired or not, based on the Hi-C map. The second category is based on the reference experiment, where the question is whether the pair is present in the reference experiment. Next, contact pairs in these categories are counted for each RNA in the target experiment and it is determined whether there are differences in the preservation of structurally associated and other contact pairs between experiments. Then the target and reference experiment are reversed. For further analysis, only cases of significance in both cases and deviations in one direction were taken (odds ratio statistics).

### 2.6. Chromatin loops

Chromatin loops were annotated using the dots function of the cooltools [32] package with default parameters (which implements an algorithm similar to HiCCUPS [17]) at a 10Kb bin size. The expected values were also calculated using cooltools. Next, the annotation of the pairs of coordinates of the loops was reduced to the linear coordinates of the loop anchors. To determine the RNAs associated with loop anchors with Fisher’s exact test, the loop anchors were expanded to 5 bins (50 Kb). We used a background model by shifting the loop anchors by 2Mb, while maintaining the regions belonging to A/B compartments to account for chromatin openness. The test was performed for each RNA separately for cis- and trans-RD-contacts, after which an adjustment for multiple testing was made using Benjamini-Hochberg procedure (threshold for q-value 0.05). For loop-level analysis and plotting, the neighborhood around the loop anchors was set to 500kb upstream and downstream.

Datasets of Hi-C mESC cells with CTCF mutated RNA binding domains were converted to cool format and preprocessed (normalized, calculated expected, annotated loops) by cooler and cooltools libraries. Loops were classified as dependent on the CTCF RNA binding domain if no loops were found in the dataset without the CTCF RNA-binding domain within 20 Kb. Otherwise, the loops were considered independent of the CTCF RNA binding domain.

### 2.7 TADs

TADs were annotated using the TopDom [36] tool, selected based on a comparative analysis [37]. The annotation was carried out on a normalized Hi-C map with a 10Kb bin size, and the window size was set to 20. We determined RNAs parental TADs by intersecting the coordinates of the RNA genes with the coordinates of the TADs (if the gene intersected with the TAD more than half its length). To demonstrate the effect of RD-scaling around the TAD in which the RNA gene is located, a neighborhood equal to the length of the TAD was selected, all TADs were reduced to a single length. The top 100 RNAs were selected according to the number of contacts in the selected region on each chromosome.

According to the position of the DNA part of the contact relative to the TAD of the gene, the contact was classified as either intra-TAD or extra-TAD. To assess the propensity of RNA to interact outside its TAD, the OutInDR parameter was used, defined as the ratio of the density of contacts outside the TAD to the density of contacts inside the TAD (InOutDR - inner/outer densities ratio). Only genes with at least 10 cis-chromosomal contacts were considered.

To search for orthologs, we used the program ortho2align [38]. The found orthologs were filtered by the Jaccard index of more than 0.5. To compare the propensity of human and mouse orthologous RNAs to leave their TAD, softer thresholds were applied during the preprocessing stage, specifically, 20% of contacts should be within peaks. Experiments for different organisms were compared (input RedC for hESC and GRID-seq and RADICL-seq for mESC).

For a set of “one-to-all” experiments, peaks of lincRNA Malat1 interactions were obtained using BaRDIC. Next, the OutInDR value was calculated.

### 3. Results

### 3.1. The contacts of many RNAs are associated with the chromatin structure

One of the main goals of this work was to determine whether there is an association between the distribution of individual RNA contacts across the genome and its three-dimensional structure. To answer this question, we proposed the procedure which is described in detail in the methods. We considered all possible pairs of contacts between certain RNA and chromatin. For each pair, we asked whether the contacts were spatially close, as defined by a Hi-C map (hereinafter referred to as paired contacts) (Figure 1). To assess the statistical significance of the observations, we used a background model by shifting the coordinates of Hi-C interactions (example in Supplementary Figure S2). The procedure was performed on 8 “all-to-all” datasets (detailed characteristics of the datasets are provided in Supplementary Table S3). For each dataset, we selected 6,000 RNAs with the largest number of peaks from the BaRDIC tool and Hi-C peaks with FDR (FitHiC2 method) 0.05. We call Hi-C significant interactions as Hi-C peaks by analogy with ChIP-seq data.

For the RADICL-seq data for the mouse embryonic stem cells (mESC) cell line, we identified 4588 RNAs whose contacts are significantly associated with chromatin structures on at least one chromosome. On the chromosome containing the RNA gene, the number of significant results (at an FDR level < 0.05) corresponding to mRNAs is approximately equal to the background number of mRNAs in the data (92.4%). On other chromosomes, results related to non-coding RNAs are overrepresented (Figure 2A). This suggests that the chromatin structure plays a role in localization around the RNA gene, regardless of its biotype. On other chromosomes it is mainly associated with contacts of non-coding RNAs, which are often involved in regulatory roles. We obtained a consistent distribution of results from other experiments (statistics for all protocols are shown on Supplementary Figure S3). Figure 2C shows paired contacts of long non-coding RNAs (mouse oligodendrocytes progenitor cells (mOPC) RADICL-seq) Malat1 and Pvt1 on chromosome 2. The distribution of paired contacts differs. Malat1 tends to interact along distant chromosomal structures, which likely represent distant interactions between the active chromatin compartments. Pvt1 interacts with more local structures (other examples on Supplementary Figure S4, Figure S5).

**Figure 2.**
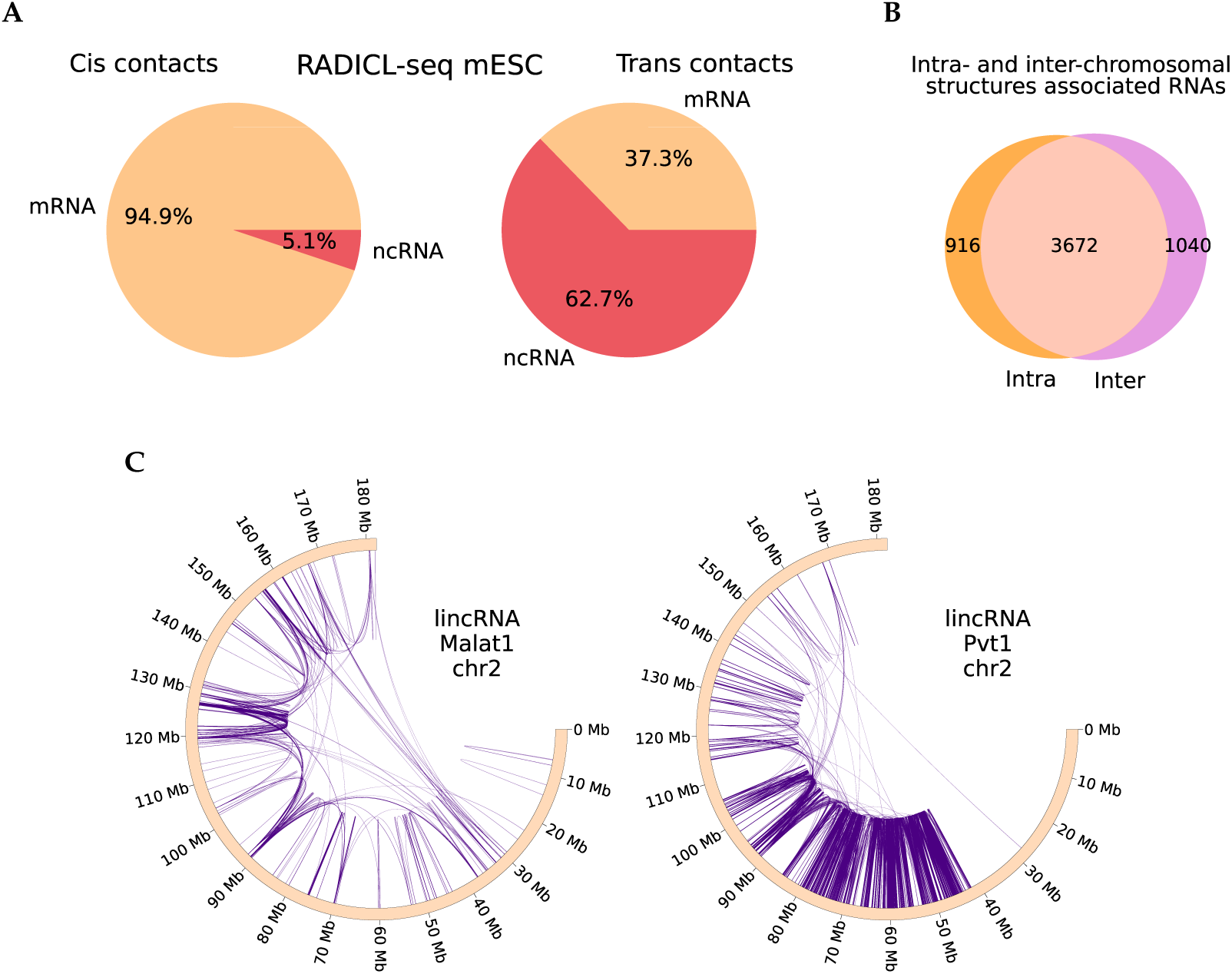
(**A**) The distribution of significant results by RNA types, on the left - on RNAs gene chromosome, on the right - on the other chromosomes, mESC RADICL-seq. (**B**) Intersections of RNAs significantly associated with the chromatin structure in the interchromosomal and intrachromosomal cases (mESC RADICL-seq). (**C**) Examples of paired contacts of long non-coding RNA of mOPC RADICL-seq data: Malat1 (chr2) on the left, Pvt1 (chr2) on the right.

Similarly, we analyzed the association of RNA-DNA interactions and chromatin structure at the interchromosomal level using FitHiC2 peaks with an FDR of 0.2. For each RNA, we considered all possible pairs of contacts between each pair of chromosomes and performed a similar calculation and statistical analysis using Hi-C interchromosomal maps. We also obtained significant results for many RNAs in both mouse (for 4712/6000 RNAs based on mESC RADICL-seq data) and human cells. Supplementary Figure S6 shows paired contacts for lncRNA Malat1 (mESC RADICL-seq) on the left. These are not separate pairs of contacts, but a “network” of many loci, which corresponds to the results of RD-SPRITE method [39]. Malat1 is involved in the formation of speckles [3], and regions associated with speckles have been shown to exhibit a tendency for interchromosomal interactions [40], which is consistent with results of this analysis. Figure 2B shows the intersections of RNAs whose contacts are associated with the chromatin structure at the intrachromosomal and interchromosomal levels. Although most RNAs show significant results in both cases, we found examples with significant paired contacts only between different chromosomes (Supplementary Table S4). This may be explained by their predominant localization at the borders of chromosomal territories. Supplementary Figure S6 on the right shows an example of interchromosomal paired contacts of such an RNA. For K562 cells, there are many instances of RNA associations with chromatin structure at the interchromosomal level (for 450 out of 6000 RNAs). However, this cell line is characterized by chromosomal rearrangements that distort both Hi-C and RNA-chromatin interactions. Many observed interchromosomal contacts and associated RNA pairs are, in fact, intrachromosomal between parts of fused chromosomes (examples on Supplementary Figure S7). This distorts the analysis, so we did not interpret the results on the K562 line.

To ensure that the obtained results were not influenced by the technical features of the “all-to-all” experiments, we analyzed data from “one-to-all” protocols for several RNAs (listed in Supplementary Table S5) from the mESC cell line. We also observed the structural association of contacts in all experiments at both the intrachromosomal and interchromosomal levels. For example, in one of the experiments for Malat1, we observed a significant enrichment of paired contacts on all chromosomes and between all chromosome pairs. Thus, with sufficient coverage, the observed relationship becomes even more pronounced.

### 3.2. Structurally associated contacts are connected with functional chromatin annotations

Next, we questioned whether we can characterize the functional role of the defined RNAs through the annotation of chromatin interactions. For K562 cells, we annotated Hi-C interactions with pairs of chromatin functional states, namely, ChromHMM and SPIN, as well as A/B compartments. We annotated the intrachromosomal paired contacts of RNAs associated with the chromatin structure and analyzed their representation in pairs of chromatin states, relative to the total number of Hi-C interactions within these states. We analyzed only pairs of states with the same name (Figures 3A,B), since loci belonging to different states interact much less intensively with each other [41,42]. For ChromHMM, in general, paired contacts are overrepresented in active chromatin (enhancers, promoters, transcription). However, there are also many cases of overrepresentation in insulators and repressed chromatin, which may be due to the involvement of a number of RNAs in the regulation of CTCF and the PRC2 complex. Paired contacts are poorly represented in heterochromatin, which is likely due to the reduced density of observed RNA contacts with inactive chromatin. For SPIN states, which characterize the association of chromatin sites with nuclear structures (Figure 3B) the propensity of individual RNAs to interact with certain states is more pronounced than in a similar plot for ChromHMM. This is likely because the authors of SPIN took into account the spatial proximity of the states of the same name based on Hi-C data, [28], which aligns with our paired contacts analysis. Within the compartments, paired contacts of most of the RNAs prevail in the active chromatin of the A compartment.

**Figure 3.**
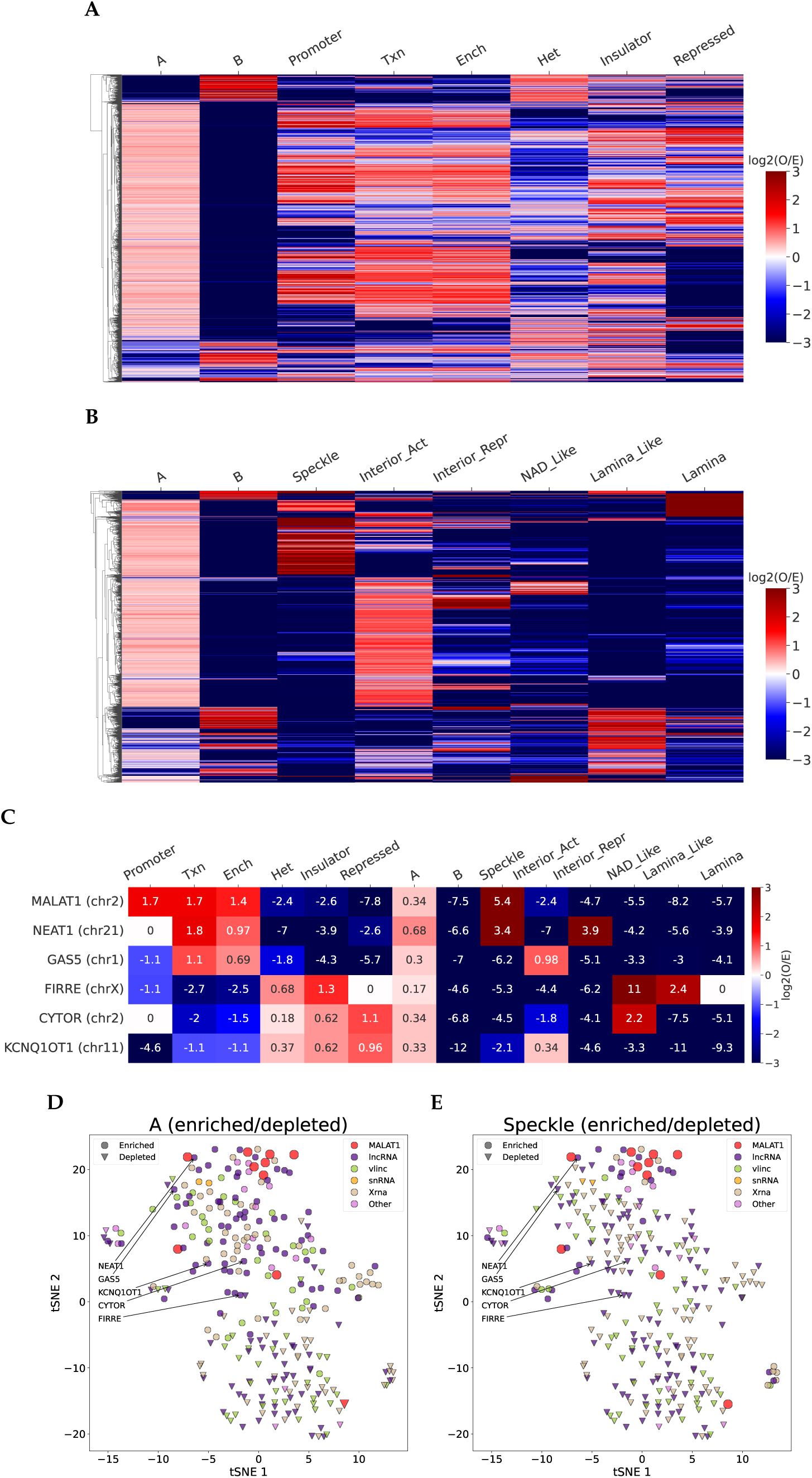
(**A,B**) Heat maps of the representation of the observed paired contacts in the states of chromatin markings relative to the expected value ((**A**) - A/B compartments + chromHMM, (**B**) - A/B compartments + SPIN states). (**C**) Heat maps for known long non-coding RNAs references. (**E,D**) Visualization of non-coding RNAs in the tSNE space based on log2 observed/expected paired contacts in all considered states, markers show enrichment/depletion of paired contacts in states pairs ((**E**) - A compartment, (**D**) - speckles).

For a comprehensive characterization of RNA functions, we combined data on all three annotations and analyzed individual long non-coding RNAs with known functions (Figure 3C). The results we observed are mostly consistent with those expected based on the literature. MALAT1 prefers to interact within the active chromatin states (ChromHMM: promoters, enhancers, transcription) on almost all chromosomes on which paired contacts have been found. In SPIN states, its paired contacts are overrepresented in speckles, which is consistent with its function in their organization (Supplementary Figure S8). The behavior of NEAT1 is generally similar to MALAT1, which is consistent with the data supporting their colocalization [43]. For the GAS5 lncRNA we primarily assume an association with active chromatin, based on the literature, [44]. We observe a similar pattern for GAS5: it preferably contacts within the active chromatin and avoids interacting with the structures of repressed and heterochromatin. As expected, paired contacts of the KCNQ1OT1 lncRNA, which is involved in imprinting the nearby locus [45], are enriched in repressed chromatin and depleted in active chromatin states. The association of FIRRE RNA with CTCF has been previously shown [6] and we observe this in the enrichment of its paired contacts in pairs of insulators. Additionally, FIRRE shows overrepresented paired contacts in nucleolus associated regions, which is consistent with the established participation of FIRRE in the colocalization of the inactivated X chromosome with the nucleolus in the mouse cells [46]. CYTOR is known to interact with the nucleolar protein nucleolin [47], and its paired contacts are overrepresented in nucleolus-associated sites.

In addition to these consistent cases, we also observe a number of contradictory results. For example, on chromosomes 12, 17 and 22, the paired contacts of MALAT1 (Supplementary Figure S8) are enriched in lamina and repressed chromatin. Also, for NEAT1, the association of paired contacts in the inactive chromatin of the SPIN annotation is observed. These inconsistent cases are in the minority and are most likely due to imperfect annotations and the noisiness of RNA-chromatin interactions data, which was not completely eliminated by the BaRDIC tool. In addition, these RNAs are highly expressed [3,43] and only a small proportion of these molecules is involved in the formation of speckles and paraspeckles.

We used the combined annotation to analyze the functions of other non-coding RNAs. Figures 3D,E show RNAs in t-SNE coordinates based on the observed-to-expected ratio of all the above-mentioned annotations. RNAs are colored by biotypes, the marker indicates cases of enrichment and depletion of paired contacts in the A compartment and speckles (other groups are shown in Supplementary Figure S9). It can be seen that RNAs are divided into several groups: NEAT1, GAS5, MALAT1, and small nuclear RNAs form a group that prefer active chromatin; KCNQ1OT1 and CYTOR are grouped based on the enrichment of paired contacts in repressed chromatin. There is a clear division of RNA into groups according to compartment preference. A group of RNAs that prefer to contact speckles is separated; MALAT1 on different chromosomes and NEAT1 form a significant part of this group.

For most of the considered RNAs with known functions, we obtained results consistent with the literature. Then we analyzed less or completely unexplored, but highly contacting RNAs for the interpretation of their possible functions based on the enrichment of paired contacts within chromatin states. We selected examples with observed-to-expected ratio > 2, adjusted p-value (chi-square) < 0.05 and the number of paired contacts on the chromosome > 1000 (the full table is given in the Supplementary Data). For promoters, we considered the very long non-coding RNA (vlincRNA) 1935_HUVEC (chr2), lncRNA AL590666.2 (chr1), ucaRNA X_11_61_b_hg38 (chr11). In addition to promoters, all of these RNAs prefer different states of active chromatin and avoid repressed or heterochromatin. One can assume that they are involved in the activation of transcription of a number of genes or in other processes within active chromatin. Examples of overrepresentation in repressed chromatin include vlincRNA 2375_K562 (chr8), ucaRNA X_10_345_b_hg38 (chr10), lncRNA LINC02476 (chr7). They may participate in the functioning of the PRC2 complex, similar to KCNQ1OT1.

### 3.3. The association of the RNA-DNA interactome and chromatin structure is conserved across experiments

It is known from the literature that the three-dimensional structure of chromatin is partially conserved across cell lines during differentiation [48]. We questioned the conservation of the RNA-DNA interactome and chromatin structure association at different levels. First, we looked at whether our results are consistent within the same cell line (mESC) across different protocols (RADICL-seq and GRID-seq) or not. Figure 4A (left) shows the intersection of RNA sets with paired contacts in each protocol, with most of the results overlapping. When comparing only non-coding RNAs (right), the intersection is also significant, but smaller, likely due to the fewer number of contacts of non-coding RNAs, which complicates the comparative analysis. Figure 4B shows similar graphs for comparing mESC and mOPC cell lines within the same protocol - RADICL-seq. The intersection is somewhat less significant than that between the results on different protocols within the same cell line, which may indicate the presence of tissue specificity in the RNA-DNA interactome association and the chromatin structure. Next, we analyzed more closely the conservativeness of paired contacts between experiments. We proposed a procedure (described in detail in the Materials and Methods section) using Fisher’s exact test. One experiment in the pair is selected as the reference, and the second as the target. The results of this for a pair of experiments RADICL-seq and GRID-seq for mESC cells are shown in Figure 4C. For most of the analyzed RNAs there is a tendency toward greater conservation of paired contacts. Since this analysis is within the same cell line, the differences are mostly explained by noise and technical biases in the experimental protocols. Thus, it can be concluded that structurally associated pairs of contacts are less noisy than those lacking spatial proximity between the same RNA’s contacts. Next, we performed a similar analysis to compare the mESC and mOPC cell lines of the RADICL-seq protocol (Fig. 4D), the results are quite similar.

**Figure 4.**
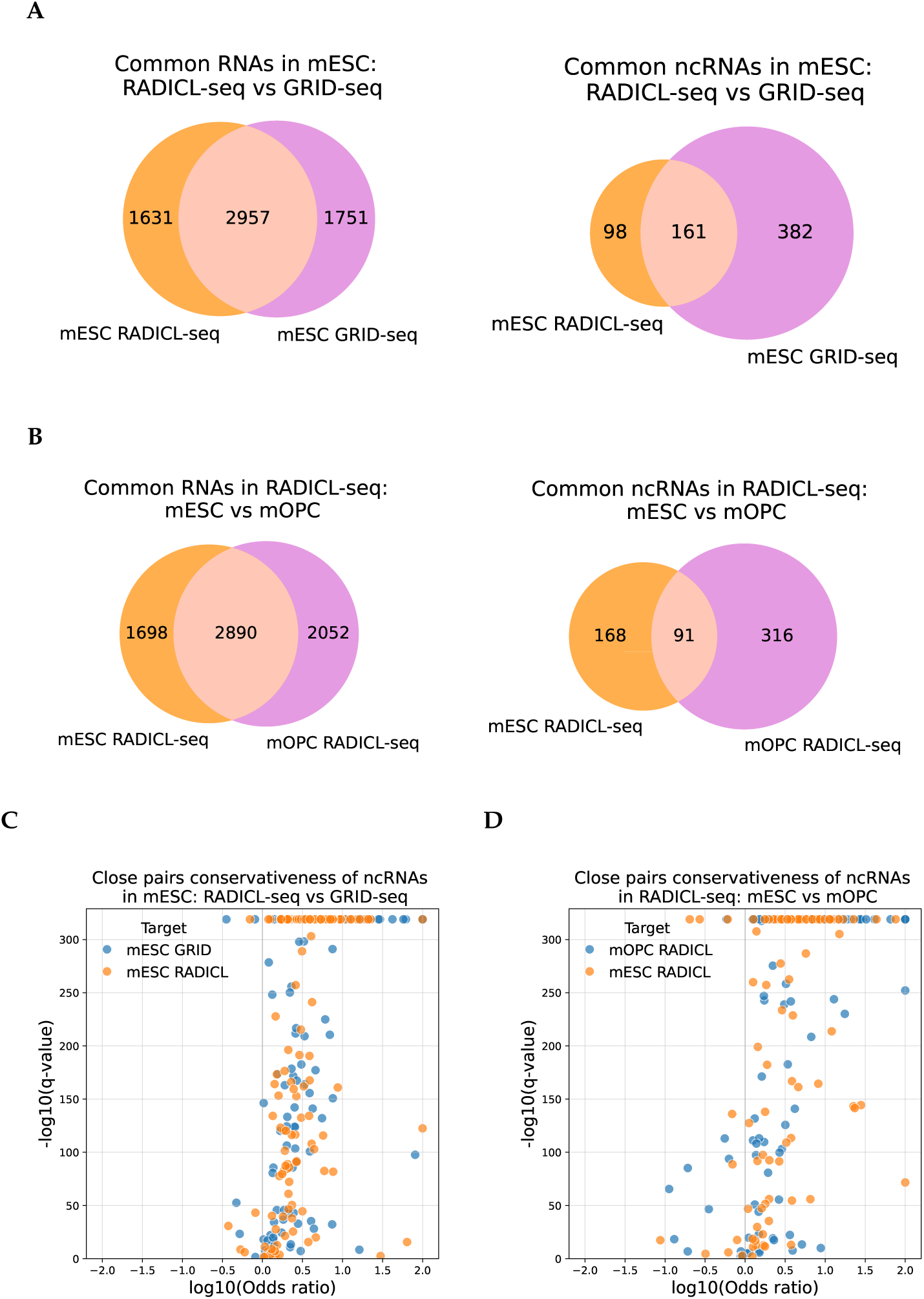
(**A,B**) Intersection of sets of RNAs associated with the chromatin structure, on the left - all RNAs, on the right - ncRNAs, (**A**) - between mESC cells based on the RADICL-seq and GRID-seq protocols, (**B**) - between mESC and mOPC cells of the RADICL-seq protocol. (**C,D**) The scatterplot of the q-values and the odds ratios of the Fisher’s exact test. (**C**) Comparison of GRID-seq and RADICL-seq protocols. (**D**) Comparison of mESC and mOPC cell lines.

Supplementary Figure S10 shows paired contacts for RNA in the mESC protocols RADICL-seq and GRID-seq with significant conservation of paired contacts across lines for RNA Malat1 and Unannotated ucaRNA X_17_3984_a_mm10. Despite the statistical significance in both cases (Fisher’s exact test: p-value adjusted: 0), similar patterns of paired contact distribution on the chromosome are much more distinguishable for ucaRNA, with many more peaks annotated for it than for Malat1 (24441 vs. 396). These two examples indicate that RNA-chromatin interaction data are sparse, and for highly contacting RNAs with many identified peaks, conservation of paired contacts is more likely to be observed. In contrast, for low-contact RNAs (Malat1 has many contacts but relatively few peaks due to its minor role in mESC [49]), this effect is underrepresented due to significant data incompleteness. Thus, this is another factor that causes underestimation of conservative cases.

The results of a similar analysis for human cell lines are shown in Supplementary Figure S11, Figure S12, Figure S13.

### 3.4. RNA contacts are associated with chromatin loops

Next, we analyzed the linear distribution of RD-contacts. The CTCF protein, known to play a key role in the formation of chromatin loops, also contains an RNA-binding domain [26]. We investigated whether RNA-chromatin interactions are associated with chromatin loops. For the analysis, we used a background model constructed by shifting the coordinates of loop anchors, while preserving their localization in A/B compartments. Supplementary Figure S14 shows the average number of RNA-DNA contacts around chromatin loop anchors (K562 Red-C). On the left random regions are shown without compartment preservation, and on the right - the A/B compartment ratio is preserved. In both cases, we observe overrepresentation of contacts at the loop anchors. However, when the distribution of loops across A/B compartments is not considered, the random model underestimates the background number of contacts. This confirms that it is crucial to account for chromatin openness in such analyses, for instance, by preserving the number of observations in each A/B compartment.

We analyzed RNA contacts on gene chromosomes, dividing them into “close” (less than 20 Mb from the gene) and “distant” categories, as well as contacts on other chromosomes. RNA preferentially contacts chromatin loops primarily near its gene. RNA is generally indifferent to distant loops (Figure 5A, mESC RADICL-seq). This is likely due to RD-scaling, as more contacts are observed near the RNA gene. Overrepresentation of contacts at loop anchors near the RNA gene is observed for most protocols we considered; however, the mOPC cell line is an exception (Supplementary Figure S15). In this case, distant contacts are also enriched at the loop anchors. Thus, we observe the tissue specificity in the interplay between RNA-DNA interactome and chromatin loops, which may indicate the functionality of this observation.

**Figure 5.**
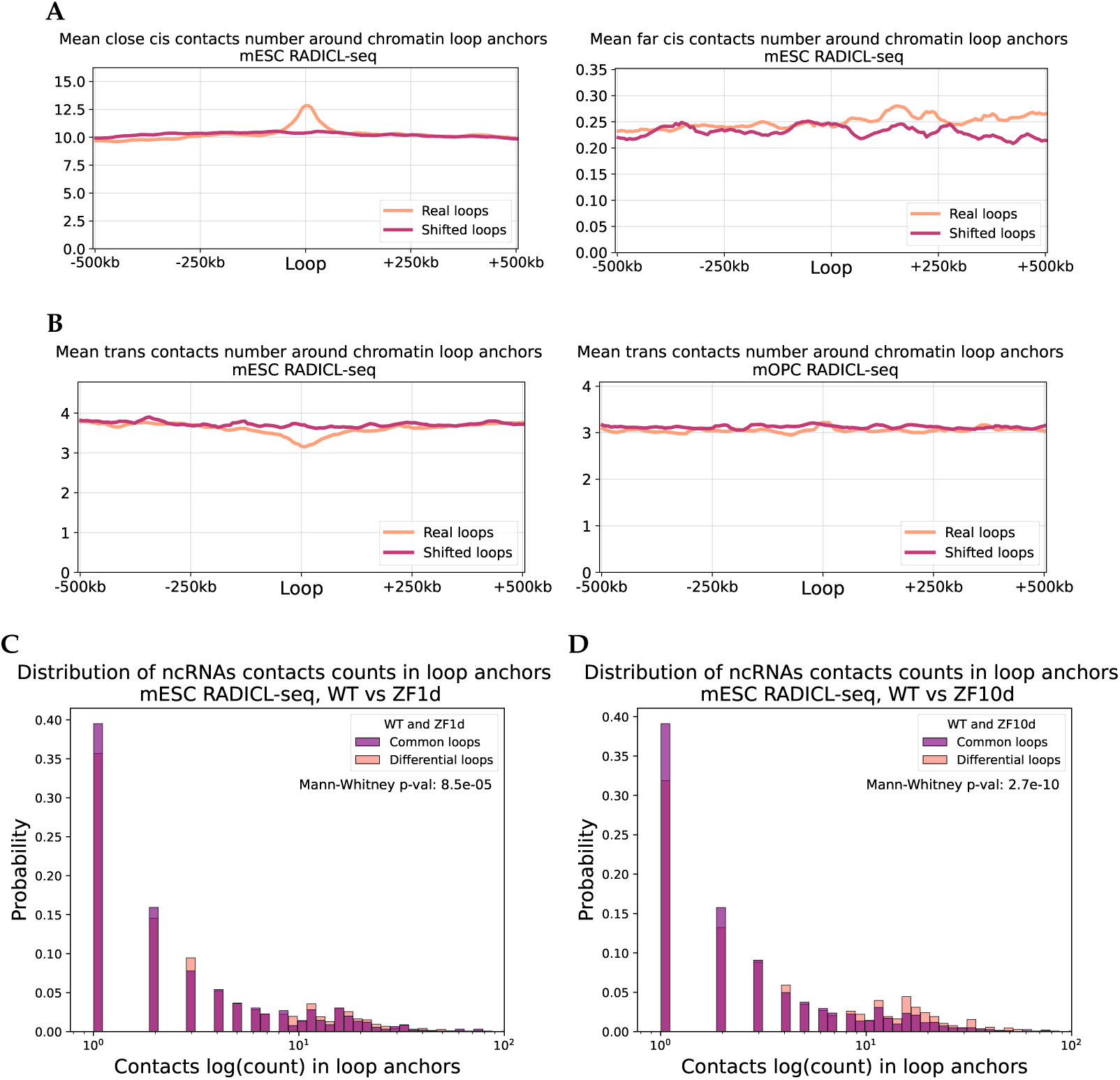
(**A**) The average number of RNA-DNA interactions in the vicinity of chromatin loop anchors, on the left - contacts closer than 20Mb from the RNA gene, on the right - further, mESC RADICL-seq. (**B**). The average number of RNA-DNA trans-interactions in the vicinity of chromatin loop anchors, on the left - mESC RADICL-seq, on the right - mOPC RADICL-seq. (**C**) Comparison of the distributions of the number of RD-contacts of the ncRNA mESC RADICL-seq in the anchors of chromatin loops. Common loops - loops anchors present in both the WT and the mutant line. Differential loops - the anchors missing in the mutant cell line (on the left is a deletion in the ZF1 domain, on the right - in the ZF10 domain).

We separately considered only the trans-contacts, Figure 5B shows the results for the RADICL-seq protocol as an example (mESC - on the left, mOPC - on the right). The result differs from long-range cis interactions: in mOPC (Supplementary Figure S15) there are no differences from the background, whereas in mESC, there is some underrepresentation of RNA contacts around the loop anchors. The results for mESC GRID-seq are consistent (Supplementary Figure S15).

We aimed to identify which RNAs contribute to this result by using Fisher’s exact test. Using RADICL-seq data for mESCs, we found significant overrepresentation of contacts in chromatin loops for 2432 RNAs. Among the known long non-coding RNAs there is Kcnq1ot1 (adjusted p-value: 1.6e-11, odds ratio: 1.5).

As demonstrated in [26], deletions in the CTCF RNA-binding domain result in the disruption of a significant number of chromatin loops. The authors introduced deletions in two domains: ZF1 and ZF10. In the case of ZF1 deletion, most loops were disrupted, whereas fewer loops were affected by ZF10 deletion. We hypothesized that RNAs might show a greater tendency to interact with loops where the presence of the CTCF RNA-binding domain is critical. We reproduced the loop annotation on three datasets, including the wild type, and obtained a consistent number of detected loops compared to the original study (WT: 5040, ZF1d: 1269, ZF10d: 4664, examples are shown in Supplementary Figure S16). Next, we divided the loops into two groups: those that persisted between the wild-type (WT) and the deletions, and those that disappeared. For these groups we analyzed the number of interactions of the RADICL-seq protocol for mESC cells. We hypothesized that if loop formation is influenced by the CTCF RNA-binding domain, we would observe more RD-interactions with these loops than with others. Supplementary Figure S17 shows the distribution of the number of contacts at the loop anchors. We observe a small but significant inverse effect: RNA-DNA interactions are more abundant in loops that do not depend on the CTCF RNA-binding domain. However, when considering only the contacts of non-coding RNAs, the expected significant effect is observed (Figure 5C, with WT vs. ZF1d on the left and WT vs. ZF10d on the right). The data for mESC GRID-seq is consistent with the RADICL-seq results, except for the lack of significance for non-coding RNAs and the ZF1 domain (Supplementary Figure S17, Figure S18).

### 3.5. RNA-DNA interactome and TADs association

We analyzed how the distribution of RNA contacts with chromatin is affected by the TAD in which the gene of this RNA is located. In earlier studies a similar analysis was carried out, but RD-scaling was not taken into account, the effect of which on RD- interactions is very significant. We selected the RNAs whose genes are located in the TADs and constructed the distribution of their contacts in the vicinity of the parental TADs. We did this both without taking RD-scaling into account (Figure 6A) and with RD-scaling considered (contacts from the peaks of the BaRDIC program, Figure 6B). We see a radical difference in the observed patterns. The result is consistent with earlier work that did not take RD-scaling into account; the intensity of interactions drops sharply at the TAD boundary, with the exception of the Red-ChIP on CTCF (see the next section, Supplementary Figure S19), where small peaks are observed at the TAD boundaries. When RD-scaling is taken into account, contacts are still overrepresented as a whole inside the TAD (shown below); however, significant peaks are observed at the boundaries of the TADs. Thus, RNA tends to interact with the boundaries of the TADs. We explain this effect by more active transcription at boundaries than inside TADs [16,50], and RNA contacts are more associated with active chromatin.

**Figure 6.**
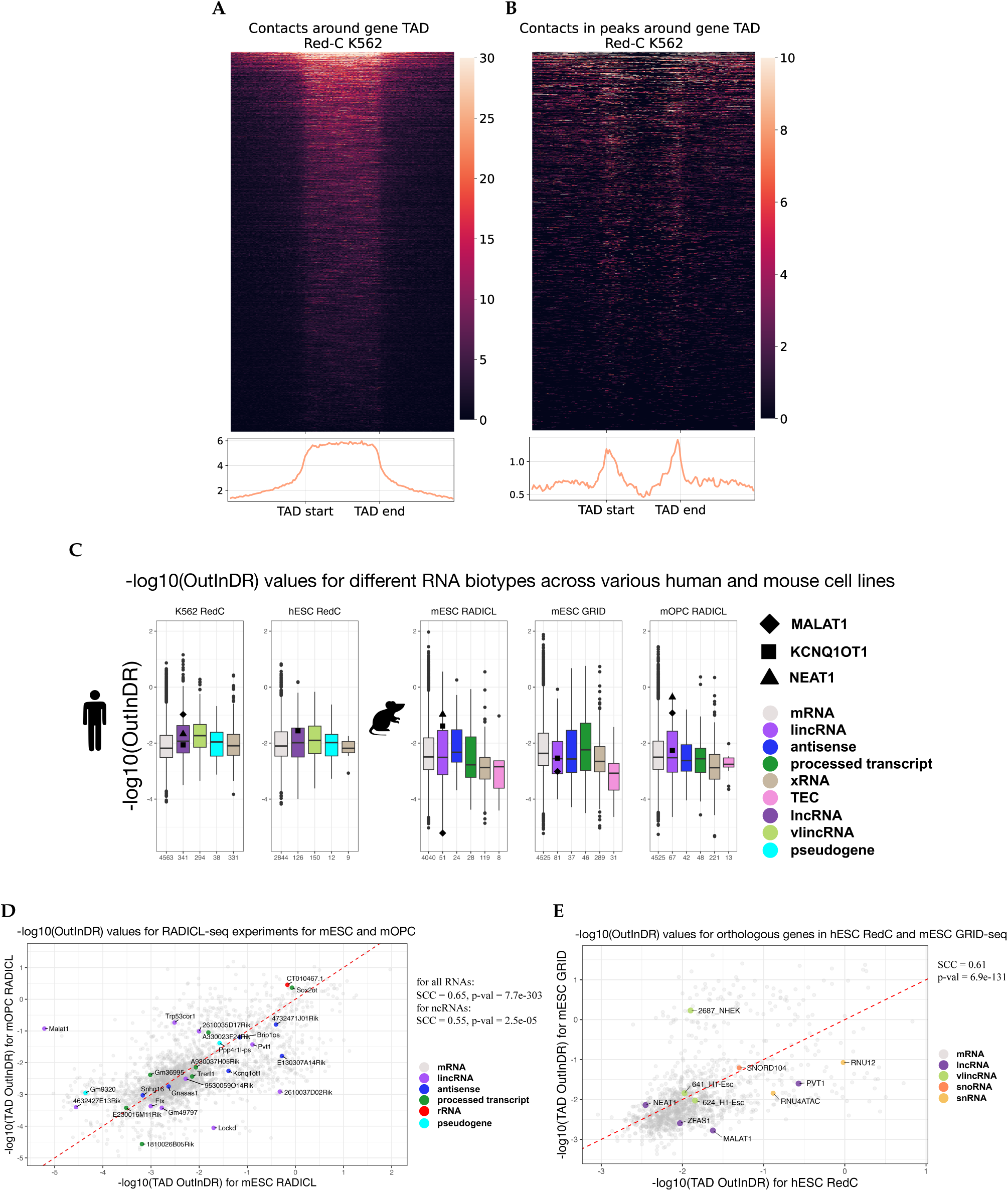
(**A**) Distribution of the full set of RD-contacts around the TADs of RNAs genes (Red-C K562). (**B**) Distribution of RD-contacts from BaRDIC peaks around the TADs of RNAs genes. (**C**) Value -log10(OutInDR) for different RNA biotypes (in different colors) for different human and mouse cell lines. The numbers presented below the graphs indicate the quantity of RNA genes included in the analysis. (**D**) Scatterplot with values -log10(OutInDR) for the RADICL-seq experiments for the mESC cell line (on the X axis) and the mOPC line (on the Y axis). (**E**) Scatterplot with values -log10(OutInDR) of orthologous genes in the hESC (X-axis) and mESC GRID-seq (Y-axis) cell lines.

We conducted a comparative analysis of RNA localization patterns on chromatin relative to their TADs for individual RNA biotypes and compared these patterns across different cell types, species and experiments. To characterize the tendency of RNA to be distributed along the chromosome, we used the ratio of peaks contact densities outside versus inside the TAD (hereinafter referred to as OutInDR). This value increases with the RNA’s tendency to leave the parental TAD and does not depend on the level of gene expression, which allows us to assess the overrepresentation of contacts outside the TAD.

We found that some RNA biotypes, such as lncRNAs and vlincRNAs, are more prone to interact outside parental TADs compared with mRNAs in K562 cells (Figure 6C). The p-values from the two-sided Mann Whitney test vary from 9.9e-7 to 2.2e-21 in experiments, as presented in the Supplementary Table S6). In the K562 and mOPC cell lines, we observe the expected pattern: KCNQ1OT1 RNA mainly functions at the locus of its gene [45], while MALAT1 and NEAT1 do not. We also observe that while MALAT1 RNA does not have a pronounced functional role in mESC [49], it is mainly localized at the site of synthesis on its chromosome. However, when MALAT1 RNA participates in the differentiation of mOPC [51], it starts to extend beyond its TAD and contact the boundaries of other TADs (p-values of the Fisher’s exact test are 1.5e-66 and 6e-30, Supplementary Figure S22). During this differentiation, NEAT1 RNA begins to cross the boundaries of its TAD and interact with other TADs (p-value of the Fisher’s exact test is 7.7e-26, Supplementary Figure S22).

To study the differences in the propensity of RNAs to contact regions outside their parental TAD in different protocols and cell lines, we analyzed the GRID-seq experiments for mESC and RADICL-seq for mESC and mOPC. We obtained a set of 1920 genes with a sufficient number of significant contacts found in all three experiments. The correlation of the OutInDR value (full set of RNAs) between different protocols within the same cell line is higher than between different cell lines for the same protocol, indicating a tissue-specific tendency of RNAs to interact regions outside their parental TADs (Figure 6D, Supplementary Figure S20). ncRNAs exhibit more conservative behavior within the same organism and more pronounced specificity in different cell lines compared to mRNAs (for mRNA GRID-seq mESC and RADICL-seq mOPC SCC = 0.88, p-val = 2.7e-17; for ncRNA, RADICL-seq mESC and RADICL-seq mOPC SCC = 0.55, p-val = 2.5e-05).

To study the conservativeness of OutInDR values across human and mouse, we compared the Red-C protocol data for hESC with the GRID-seq and RADICL-seq data for mESC. A list of 1283 orthologous genes was obtained. Then we compared the OutInDR value for each gene in the hESC and mESC cell lines (Figure 6E, Supplementary Figure S21). We observed conservativeness in the tendency of orthologous human and mouse RNA to contact areas outside their parental TADs (for GRID-seq: SCC = 0.61, p-value = 6.9e-131, for RADICL-seq: SCC = 0.59, p-value = 6.2e-120).

We compared the OutInDR values of the lincRNA Malat1 between the “all-to-all” and “one-to-all” experimental data. A comparison of the “all-to-all” data of the GRID-seq experiment for mESC and the data of 7 “one-to-all” experiments (Supplementary Figure S23) showed that Malat1 exhibits a greater tendency to leave its TAD in the “one-to-all” experiments compared to the “all-to-all” data (the value is log10(OutInDR) in the experiment GRID-seq for mESC was -2.77, the median log10(OutInDR) value in the “one-to-all” experiments is -1.07). In an experiment where cells were treated with a transcription elongation inhibitor [52], there was a significant decrease in Malat1’s ability to interact to areas outside its TAD, which is expected due to Malat1’s role in speckles.

### Analysis of RNA-DNA-protein interactions

An experimental protocol, Red-ChIP, has been proposed, which allows researchers to fix RNA-DNA interactions with an additional step of immunoprecipitation for a specific protein [53]. In this study, 4 datasets were obtained: one from K562 cells with immunoprecipitation for CTCF protein, one from hESC cells with immunoprecipitation for EZH2 (a component of the polycomb repressive complex), and two additional Red-C datasets (as input) for the same cell lines. We repeated the above analysis to determine the association of RNA with chromatin structure and identified many significant RNAs, both for cis- and trans-RD-interactions (Supplementary Figure S3).

We annotated paired contacts with pairs of ChromHMM states. The insulator state is similar to CTCF peaks, while the repressed chromatin state is characterized by peaks of PRC2 complex proteins. The number of RNAs whose paired contacts are enriched or depleted in the insulator states in both the input and Red-ChIP datasets (on CTCF) is shown in Figure 7A (similar for RedChIP on EZH2 in the Repressed state in Figure 7B). We observe significantly greater representation of paired contacts in the corresponding chromatin states for Red-ChIP. At the same time, for example, paired contacts in RedChIP on EZH2 are depleted in the state of active transcription (Supplementary Figure S24).

**Figure 7.**
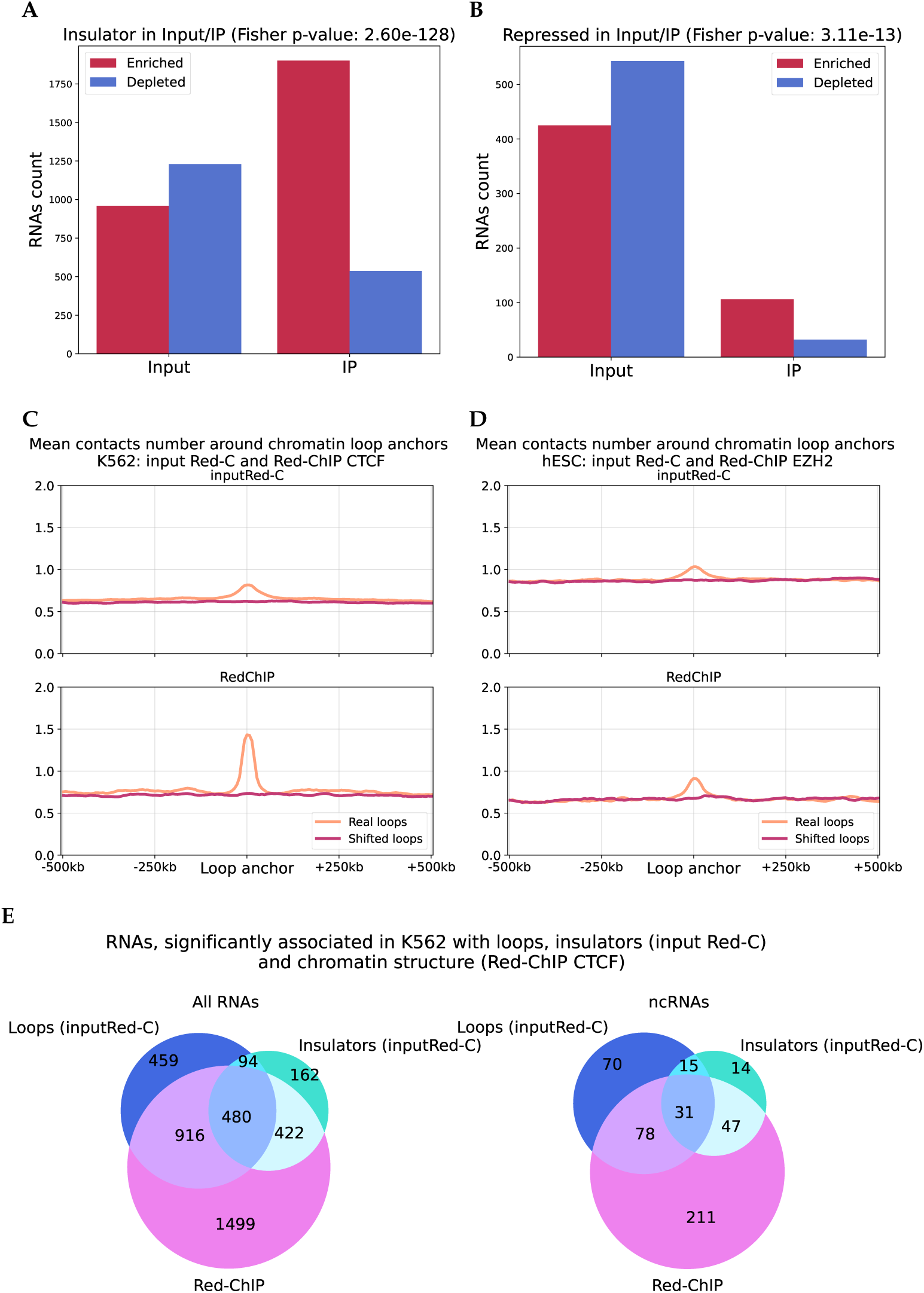
(**A,B**) The number of cases of significant enrichment and depletion of paired contacts in pairs of states in Red-C and Red-ChIP. (**C**) Comparison of the representation of contacts within the chromatin loop anchors (K562), on the left - Red-ChIP on CTCF, on the right - Red-C (the number of contacts is normalized to the number of significant contacts in the experiment). (**D**) Comparison of the representation of contacts within the chromatin loop anchors (hESC), on the left - Red-ChIP on EZH2, on the right - Red-C (the number of contacts is normalized to the number of contacts from peaks in the experiment). (**E**) The intersection of RNAs significantly associated with loops (input Red-C data, K562), with the chromatin structure (Red-ChIP), and whose paired contacts are associated with insulators (input-Red-C).

We also studied the protocol differences in the context of chromatin loops. For contacts fixed via CTCF, we observe a much clearer and higher peak in the loop anchors compared to conventional Red-C (Figure 7C), which is completely consistent with the role of CTCF in the organization of chromatin loops. For immunoprecipitation via EZH2, no such prominent peak is observed (Figure 7D), since this protein is not involved in the formation of chromatin loops. Additionally, for Red-ChIP on CTCF, we observe an association with chromatin loops not only for contacts near the gene, but also of distant contacts and trans-contacts, in contrast to the data from Red-C (Supplementary Figure S15).

To isolate RNAs consistently associated with loops and CTCF, we compared the results of the analysis of paired contact enrichment with insulator states (input Red-C), RNA association with loop anchors (input Red-C), and RNAs associated with the chromatin structure in the Red-ChIP protocol (CTCF, K562 cell line). We found a significant number of intersections (Figure 7E). We selected those RNAs whose results intersect on specific chromosomes and that also overlap with RNAs identified from another Red-C experiment (Supplementary Figure S25): LINC01184 (DNA parts: chr5), GARS1-DT (chr7), AC073529.1 (chrX), 1840_HUVEC (chr7), 1366_HepG2 (chr7). These RNAs are most likely to be involved in the regulation of CTCF.

## 4. Discussion

For a significant number of non-coding RNAs, their role in chromatin regulatory processes, structural organization, and interactions with specific proteins is well established. The number of datasets of RNA-DNA interactions is constantly growing, expanding the possibilities for analyzing the functions of chromatin-associated RNAs. In the HiChIRP [20] protocol, the first joint experimental study of RNA-chromatin interactions and the spatial structure of chromosomes for several RNAs in the “one-to-all” mode was performed. In this work, we applied a similar in-silico analysis for genome-wide “all-to-all” data. We have identified RNAs associated with chromatin structure; moreover, we have shown that this property appears to be universal for many RNAs, which allows us to conclude that RNAs are localized along the genome in a manner that aligns with the chromosomal structure, rather than linearly. Furthermore, RNA contacts are not independent events; their distribution depends on the pairwise proximity of the corresponding genome loci. We observed this effect both within the same chromosome and between different chromosomes. We analyzed this effect on 8 sets of experimental data from different protocols, cell lines and organisms, showing consistent results. Moreover, we have shown that there is a conservation between different experiments. Within a single cell line, most RNAs tend to interact with the same chromosomal structures, which indicates that these pairs of interactions are less noisy. Conservation is also observed between different cell lines.

We analyzed the chromosomal structures with which individual RNAs are associated in the context of functional chromatin annotations. We found that the preferences of RNAs for specific chromosomal structures differ. For RNAs with known functions, we observed consistent results: MALAT1 preferentially interacts within speckles and active chromatin, while KCNQ1OT1 is enriched in repressed chromatin structures.

All of this highlights the importance of considering chromatin structure in the analysis of RNA-chromatin interactions. Although early studies focused on the influence of chromatin openness and the effect of RD-scaling on these data, the role of spatial factors in the distribution of RNA-chromatin interactions has not been comprehensively studied until now. We have demonstrated its significant role in different examples.

We also analyzed how chromatin loops are related to RNA-chromatin interactions. Our analysis shows that, in general, RNAs tend to contact the anchors of the loops closest to the RNA gene, and this pattern is observed for all data sets. It is further confirmed by the fact that in the RedChIP experiment data, which includes an immunoprecipitation stage with CTCF, this effect is especially obvious. Additionally, we analyzed Hi-C data for a mESC cell line with a mutated CTCF RNA-binding domain. We assumed that, since many chromatin loops disappear, we may see differences in the distributions of RD-contacts with these loops. This hypothesis has not been confirmed for a full RNA set, but such an effect has been shown for ncRNAs.

We also conducted a comprehensive analysis of the relationship between RNA-chromatin interactions and TADs. First, we showed that it is critically important for this analysis to take into account RD-scaling, since this shows the preference of RNA for interacting with the boundaries of TADs. We showed that individual RNA biotypes exhibit different localization patterns on chromatin relative to their TAD, and found that these patterns are mostly preserved across different protocols, cell lines, and even organisms (human and mouse). However, we also observed some degree of cellular specificity in this behavior, suggesting that although the tendency of RNA to localize beyond its TAD is conserved, it may also be influenced by tissue-specific factors. We found a change in the localization pattern of MALAT1 RNA as it differentiated from mouse embryonic stem cells into oligodendrocyte progenitor cells. In mESC, it localizes at the site of synthesis, but as it acquires a functional role in mOPC, it begins to extend beyond its parental TAD and contact the boundaries of other TADs located in more distant parts of the genome. Analysis of orthologous genes in human and mouse embryonic stem cells revealed a significant positive correlation, indicating that the tendency of RNAs to bind to regions outside their parental TADs is partially preserved between species.

We also found previously uncharacterized non-coding RNAs that may be associated with chromatin structure.

## Author Contributions

Conceptualization, A.M.; methodology, D.Z., A.Z. and A.M.; software, D.Z. and A.T.; validation, D.Z., A.Z., and A.M.; formal analysis, D.Z. and A.T.; investigation, D.Z. and A.T.; resources, A.Z.; data curation, D.Z. and A.Z.; writing—original draft preparation, D.Z. and A.T.; writing—review and editing, D.Z., A.Z. and A.M.; visualization, D.Z. and A.T.; supervision, A.Z. and A.M.; project administration, A.Z. and A.M.; funding acquisition, A.M. All authors have read and agreed to the published version of the manuscript.

## Institutional Review Board Statement

Not applicable

## Data Availability Statement

Hi-C data were taken from the 4DNucleome database (K562: 4DNESI7DEJTM, hESC: 4DNES2M5JIGV, mESC: 4DNESDXUWBD9, mOPC: 4DNESJ9SIAV5) and GEO: GSE125595. RNA-DNA interactomes were taken from RNAChromDB (ID for “one-to-all” experiments: 36, 49, 52, 55, 58, 92, 93, 94, 156, 198).

## Acknowledgments

The authors thank Alexey Shkolikov for their valuable comments on manuscript.

## Abbreviations

The following abbreviations are used in this manuscript:

RD: RNA-DNA
lncRNA: Long noncoding RNA (human)
vlincRNA: Very long intergenic noncoding RNA (human)
lincRNA: Long intergenic noncoding RNA (mouse)
X RNA,: ucaRNA Unannotated chromatin associated
RNA: snRNA Small nuclear RNA
snoRNA: Small nucleolar RNA
TAD: Topologically associated domain
mESC: Mouse embryonic stem cells
mOPC: Mouse oligodendrocytes progenitor cells
hESC: Human embryonic stem cells
OutInDR: Contacts densities ratios outside versus inside the TAD

