## Supplementary Note for "Joint analysis of RNA-DNA and DNA-DNA interactomes reveals their strong association"

We varied the shift value (0.5, 2, 10 megabases) and got fairly consistent results (Supplementary Figure S1). We settled on 2 megabases, since this value exceeds the size of most of the TADs (data in the literature differs greatly depending on the algorithm, data coverage and cell line: 880 Kb for mouse [1], 185 Kb for human [2]; using the TopDom[3] we got 290 Kb for K562 and 180 Kb for mESC). This allows us to assume a sufficient level of independence between the real and background models. When shifting the coordinates of the Hi-C peaks, we kept them belonging to A/B compartments, in order to preserve the plaid pattern on the Hi-C map and take into account the openness of chromatin (the importance of this is demonstrated in the results section about chromatin loops, Supplementary Figure S14).

When setting a universal threshold for peaks q-values, the total number of peaks in different data sets differs greatly (in some cases by orders of magnitude). This can be explained by the different reads coverage and differences in the experimental procedure, which lead to different data quality, which causes differences in the binomial assessment of the significance of interactions. A similar problem occurs when analyzing ChIP-seq [4] data, where it is often solved by analyzing the reproducibility level of peaks between replicas, instead of setting a universal threshold for q-value. To smooth out these technical differences, we varied the thresholds by the q-value of peaks for different data sets, based on the assumption that the data is very noisy and the proportion of specific interactions should not be large. We have chosen a threshold of 10% of contacts falling into peaks.
