## Supplementary Information for "Joint analysis of RNA-DNA and DNA-DNA interactomes reveals their strong association"

S12. The intersection of the sets of RNAs associated with the chromatin structure between the protocols of the K562 and hESC cell lines, the Red-C (input) protocol, on the left - all RNAs, on the right - non-coding RNAs. 17
S13. The relationship of the q-values and the odds ratios of the Fisher's exact test. On the left - comparison of Red-C and Red-C (input) experiments, on the right comparison of K562 and hESC cell lines. . . . . 18
S14. The average number of RNA-DNA (K562 Red-C) interactions in the vicinity of chromatin loop anchors, on the left the background model is random shift, on the right the background model is shift, taking into account belonging to the A/B compartment. . . . . 18
S15. The average number of RNA-DNA interactions in the vicinity of chromatin loop anchors (continued on the next page). . . . . 19
S15. The average number of RNA-DNA interactions in the vicinity of chromatin loops anchors. . . . . 20
S16. Examples of annotated loops on Hi-C maps in different conditions, on the left - WT, in the center - deletion in ZF10, on the right - deletion in ZF1 (chr17:30330000:32900000). . . . . 21
S17. Comparison of the distribution of the number of RD-contacts in mESC within the anchors of chromatin loops. Common loops are found in both WT and mutant lines. Differential loops are absent in the mutant cell line (on the left is a deletion in the ZF1 domain, on the right is in the ZF10 domain, the upper row is RADICL-seq, the lower one is GRID-seq). . . . . 21
S18. Comparison of the distribution of the number of ncRNAs RD-contacts in mESC GRID-seq within the anchors of chromatin loops. Common loops are found in both WT and mutant lines. Differential loops are absent in the mutant cell line (on the left is a deletion in the ZF1 domain, on the right is in the ZF10 domain. 22
S19. Left - distribution of the full set of RD-contacts around the TADs of RNAs genes (Red-C K562), right - distribution of RD-contacts from BaRDIC peaks around the TADs of RNAs genes (continued on the next page). . . . . 23
S19. Left - distribution of the full set of RD-contacts around the TADs of RNAs genes (Red-C K562), right - distribution of RD-contacts from BaRDIC peaks around the TADs of RNAs genes (continued on the next page). . . . . 24

S19. Left - distribution of the full set of RD-contacts around the TADs of RNAs genes (Red-C K562), right - distribution of RD-contacts from BaRDIC peaks around the TADs of RNAs genes (continued on the next page). . . . . . 25
S19. Left - distribution of the full set of RD-contacts around the TADs of RNAs genes (Red-C K562), right - distribution of RD-contacts from BaRDIC peaks around the TADs of RNAs genes. . . . . . 26
S20. Scatterplot with values -log10(OutInDR) for the RADICL-seq experiment on the X axis and GRID-seq on the Y axis for the mESC cell line. . . . . . 27
S21. Scatterplot with values -log10(OutInDR) of orthologous genes in the hESC (X-axis) and mESC RADICL-seq (Y-axis) cell lines. . . . . . 28
S22. RD-contacts distribution in TADs and boundaries in different cell lines. Top row - MALAT1 contacts, bottom row - NEAT1. . . . . . 28
S23. Comparison of the -log10(OutInDR) value for lincRNA Malat1 in "one-to-all" and "all-to-all" experiments for mESC GRID-seq. . . . . . 29
S24. The number of cases of significant overrepresentation and underrepresentation of paired contacts in pairs of the active transcription state in Red-C and RedChIP. . . . . . 29
S25. The intersection of RNAs which are significantly associated with loops (Red-C data), with the chromatin structure (Red-ChIP), and whose paired contacts are associated with insulators (Red-C). . . . . . 30

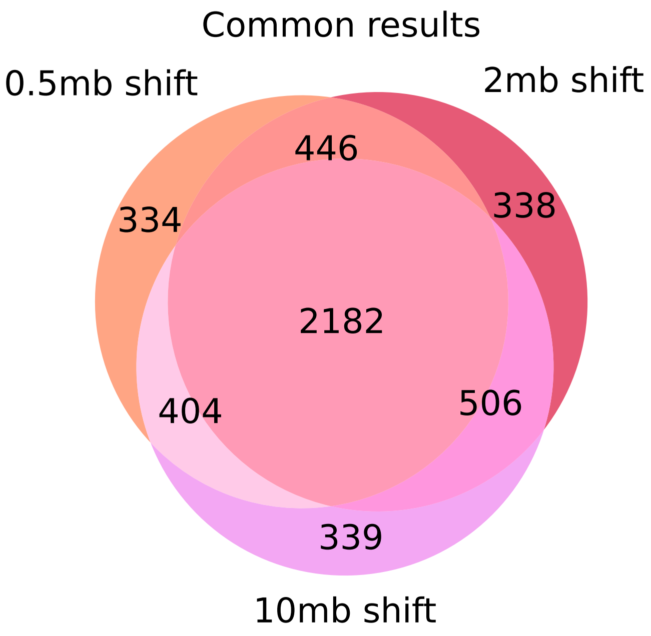

**Figure S1.** The intersection of the results of significant association of RNA-DNA interactions and chromatin structure depending on the shift (K562 Red-C). The numbers of different results are close between the shift values, which indicates that the magnitude of the shift does not affect the degree of difference between the real model and the background one.

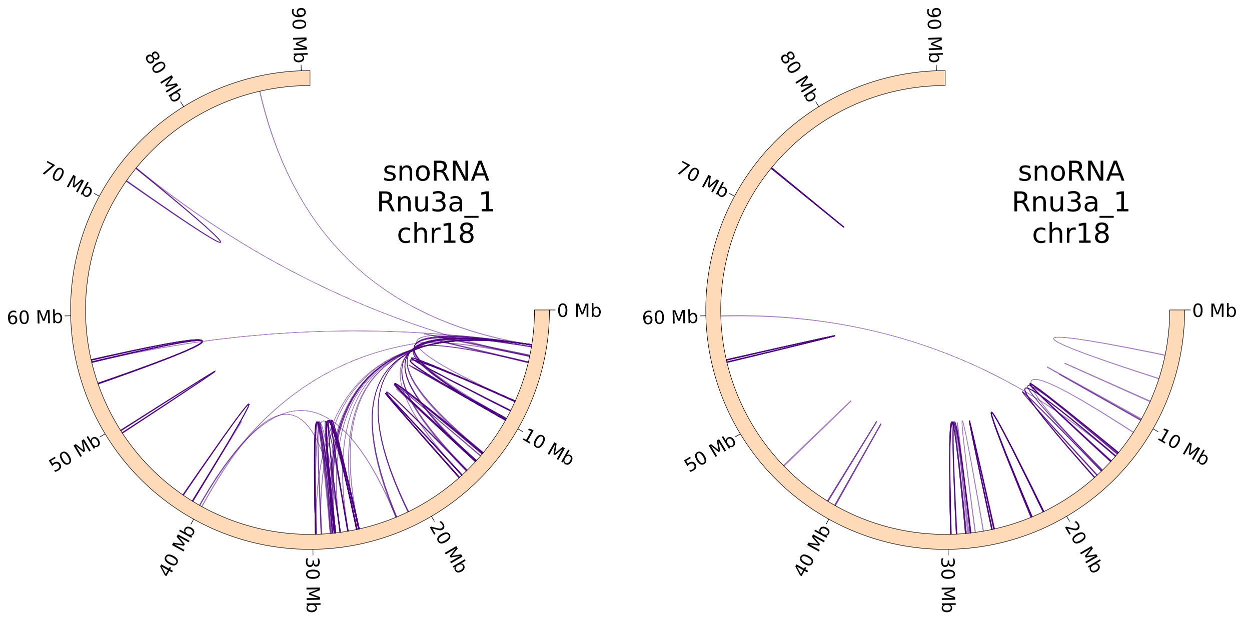

**Figure S2.** Paired Rnu3a RNA contacts (mESC RADICL-seq) based on real (left) and background (right) Hi-C maps. The number of paired contacts in the background model is significantly lower (p-value adjusted: 2.3e-95, odds ratio: 4.5), which clearly confirms the validity of our procedure.

**Table S1.** ChromHMM states grouping [1].

| Grouped state | ChromHMM state |
|---|---|
| Promoter | 1_Active_Promoter<br>2_Weak_Promoter<br>3_Poised_Promoter |
| Ench | 4_Strong_Enhancer<br>5_Strong_Enhancer<br>6_Weak_Enhancer<br>7_Weak_Enhancer |
| Insulator | 8_Insulator |
| Txn | 9_Txn_Transition<br>10_Txn_Elongation<br>11_Weak_Txn |
| Repressed | 12_Repressed |
| Het | 13_Heterochrom/lo<br>14_Repetitive/CNV<br>15_Repetitive/CNV |

**Table S2.** SPIN states grouping [2].

| Grouped state | SPIN state |
|---|---|
| Speckles | Speckles |
| Interior_Act | Interior_Act1<br>Interior_Act2<br>Interior_Act3 |
| Interior_Repr | Interior_Repr1 |
| NAD_like | Interior_Repr2 |
| Lamina_Like | Lamina_Like<br>Near_Lm1<br>Near_Lm2 |
| Lamina | Lamina |

**Table S3.** Characteristics of RD-protocols.

| Protocol | Paper | Cell line | Interactions number | Interactions set | BaRDIC peaks count | FDR threshold |
|---|---|---|---|---|---|---|
| Red-C | [3] | K562 | 43603491 | All<br>Cis | 663396<br>91661 | 0.203<br>0.11 |
| Red-C (input) | [4] | K562 | 29288865 | All<br>Cis | 942379<br>59040 | 0.015<br>0.003 |
| Red-C (input) | [4] | hESC | 16158007 | All<br>Cis | 238727<br>45194 | 0.119<br>0.055 |
| Red-ChIP | [4] | K562 | 19323288 | All<br>Cis | 645219<br>40878 | 0.096<br>0.012 |
| Red-ChIP | [4] | hESC | 4041758 | All<br>Cis | 75271<br>21843 | 0.048<br>0.048 |
| RADICL-seq | [5] | mESC | 47319620 | All<br>Cis | 564435<br>54605 | 0.171<br>0.001 |
| RADICL-seq | [5] | mOPC | 45974349 | All<br>Cis | 522858<br>60127 | 0.187<br>0.007 |
| GRID-seq | [6] | mESC | 62398550 | All<br>Cis | 443485<br>73272 | 0.02<br>0.00015 |

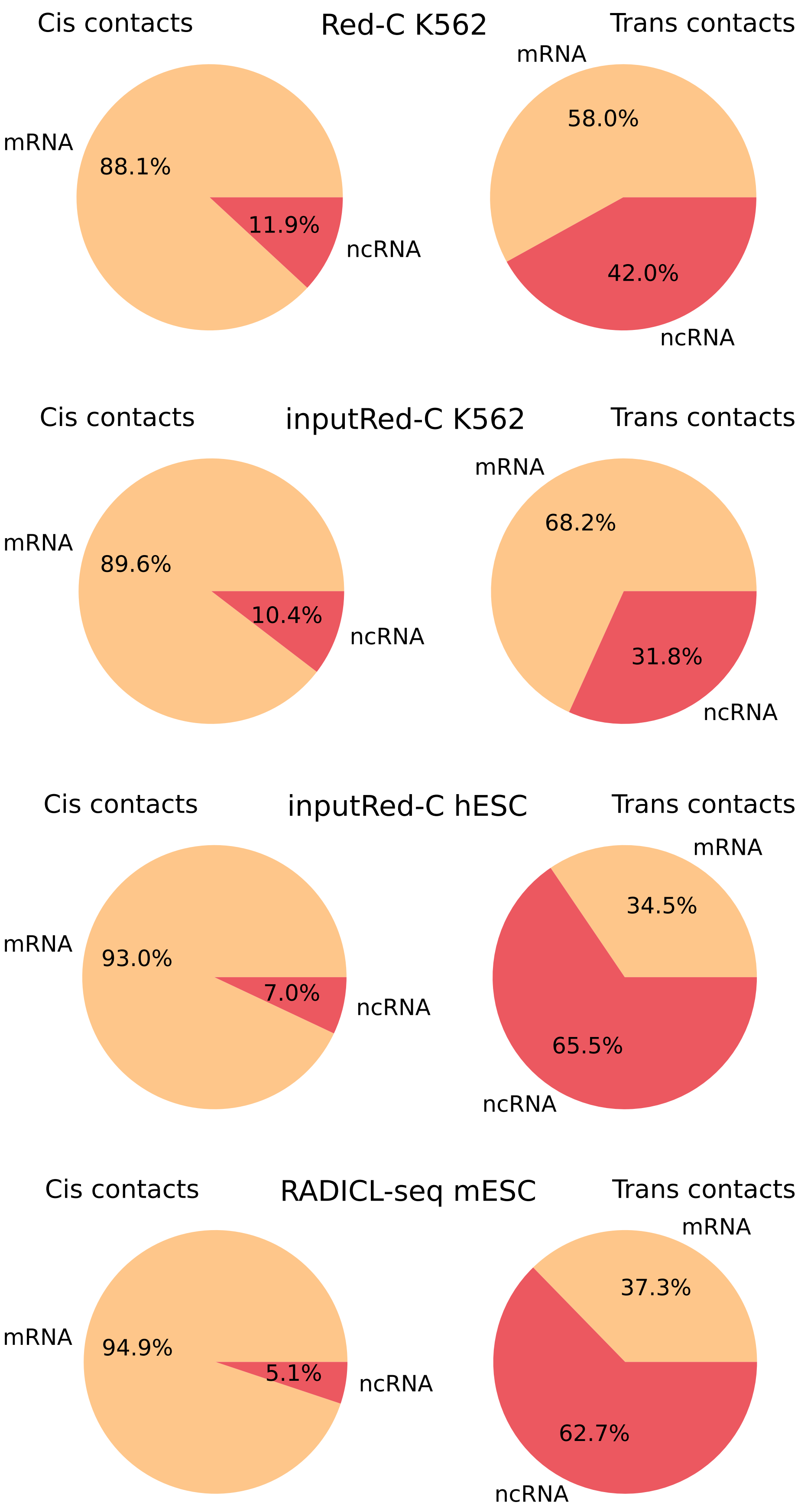

**Figure S3.** Distribution of significant results by RNA types (continued on the next page).

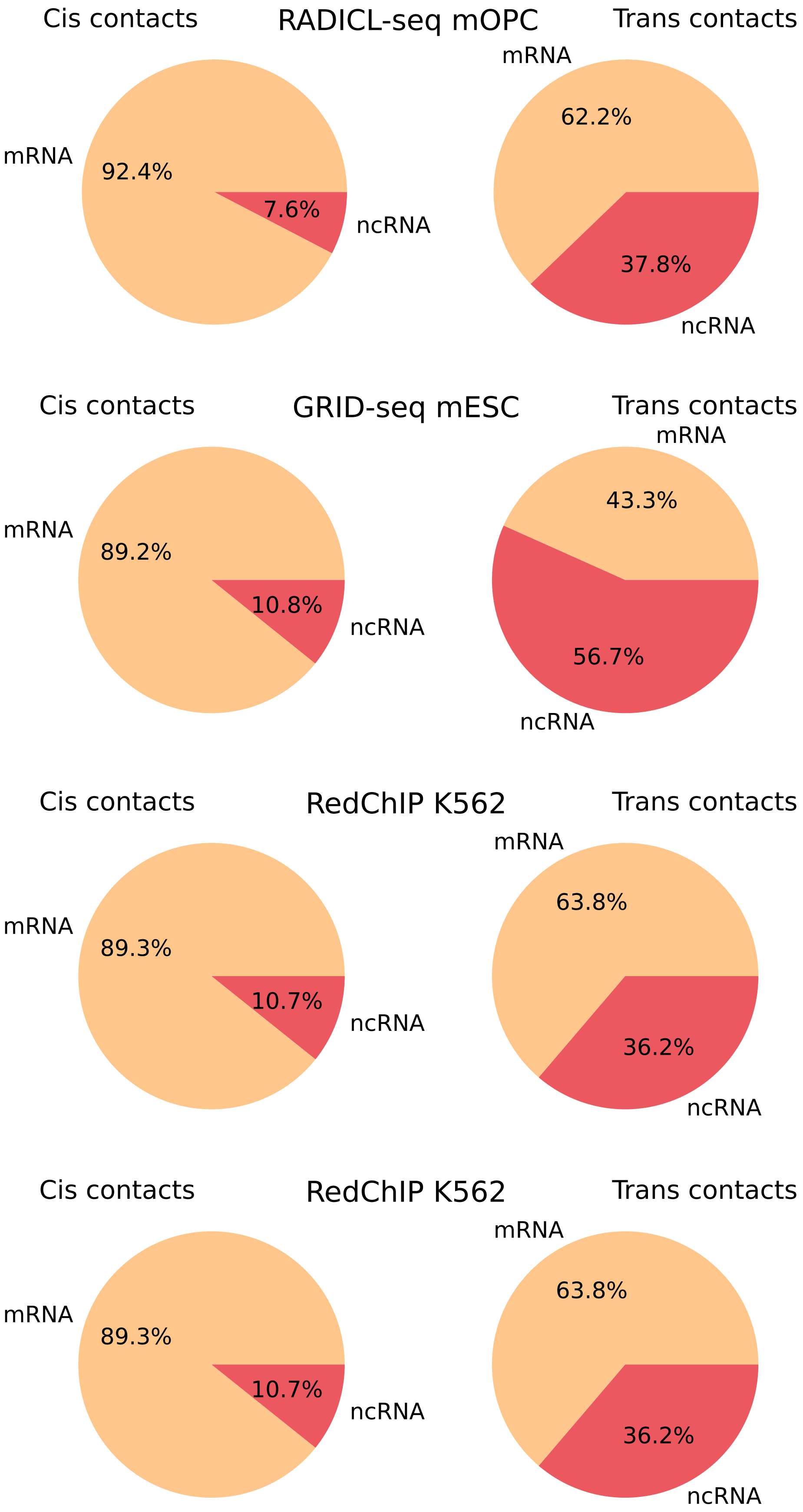

**Figure S3.** Distribution of significant results by RNA types

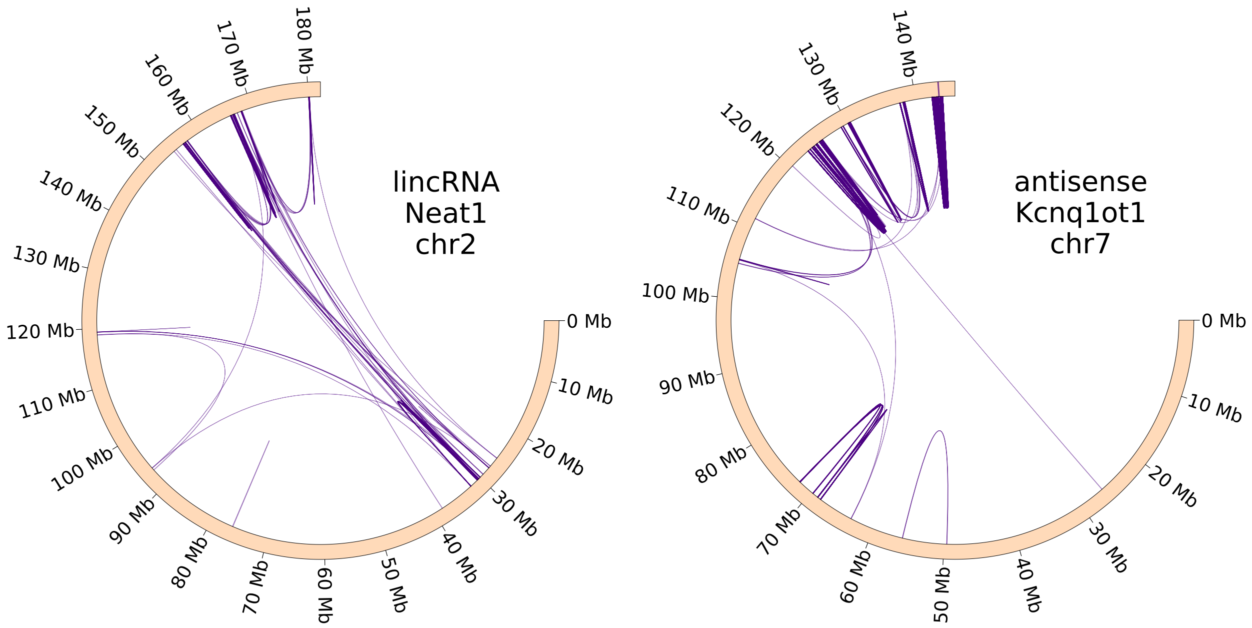

**Figure S4.** Examples of paired contacts of lncRNA of mOPC RADICL-seq data: Malat1 (chr15) on the left, Pvt1 (chr15) on the right.

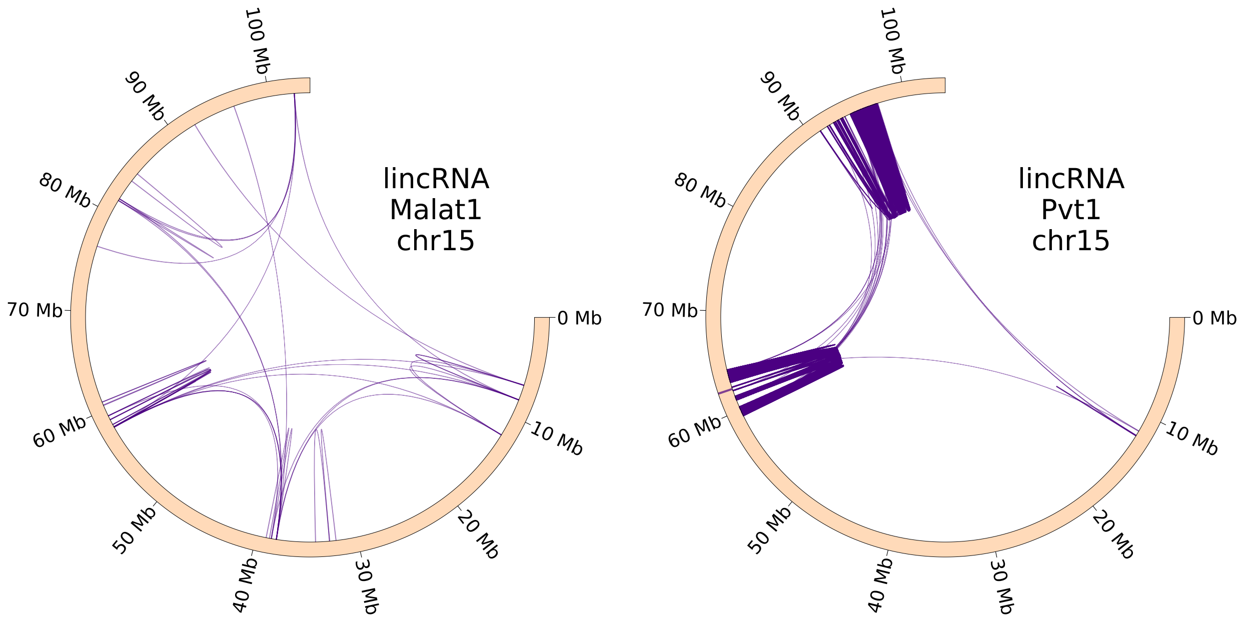

**Figure S5.** Examples of paired contacts of lncRNA of mOPC RADICL-seq data: Neat 1 (chr2) on the left, Kcnq1ot1 (chr7) on the right.

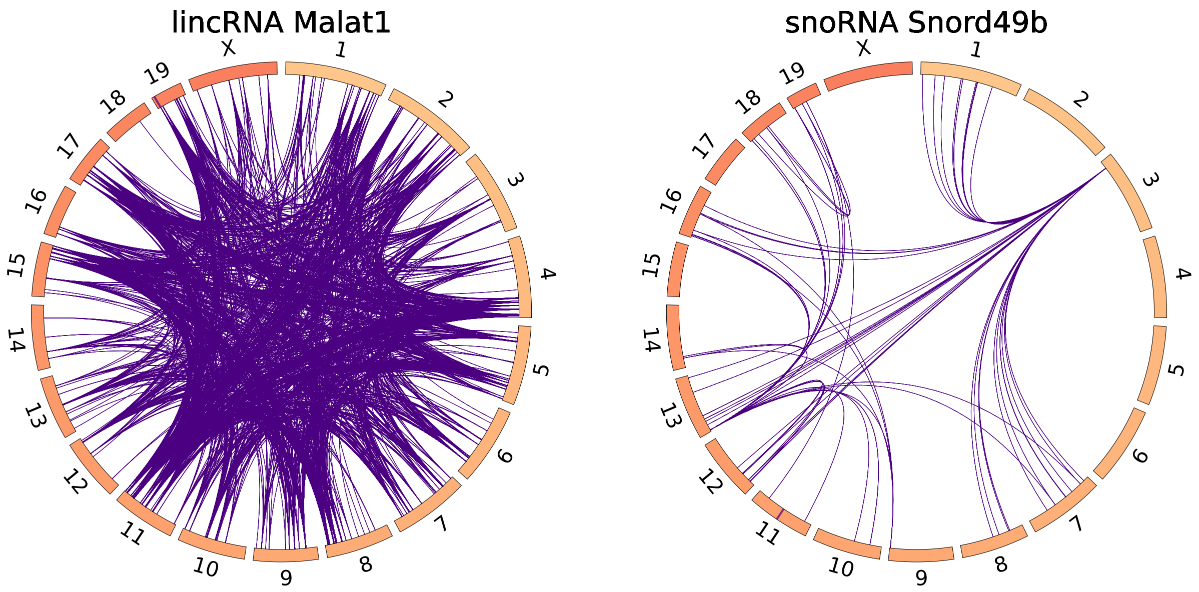

**Figure S6.** Examples of interchromosomal paired contacts of long non-coding RNAs (mESC RADICL-seq).

**Table S4.** Non-coding RNAs (mESC RADICL-seq) associated with chromatin structure only at the interchromosomal level. RNAs associated with more than 4 pairs of chromosomes are shown.

| RNA | RNA type | Number of chromosome pairs |
|---|---|---|
| Snord49b | snoRNA | 19 |
| Mirg | lincRNA | 12 |
| Snord47 | snoRNA | 11 |
| Lncpint | lincRNA | 9 |
| LSU-rRNA_Hsa_252 | rRNA | 9 |
| 1600020E01Rik | antisense | 8 |
| Gm50050 | lincRNA | 7 |
| Pisd-ps2 | transcribed_unprocessed_pseudogene | 7 |
| X_4_13396_c_mm10 | Xrna | 7 |
| Gm27533 | misc_RNA | 6 |
| Snhg1 | processed_transcript | 6 |
| Snord43 | miRNA | 6 |
| SSU-rRNA_Hsa_21 | rRNA | 6 |
| B830012L14Rik | lincRNA | 5 |
| Dancr | processed_transcript | 5 |
| Zfas1 | processed_transcript | 5 |

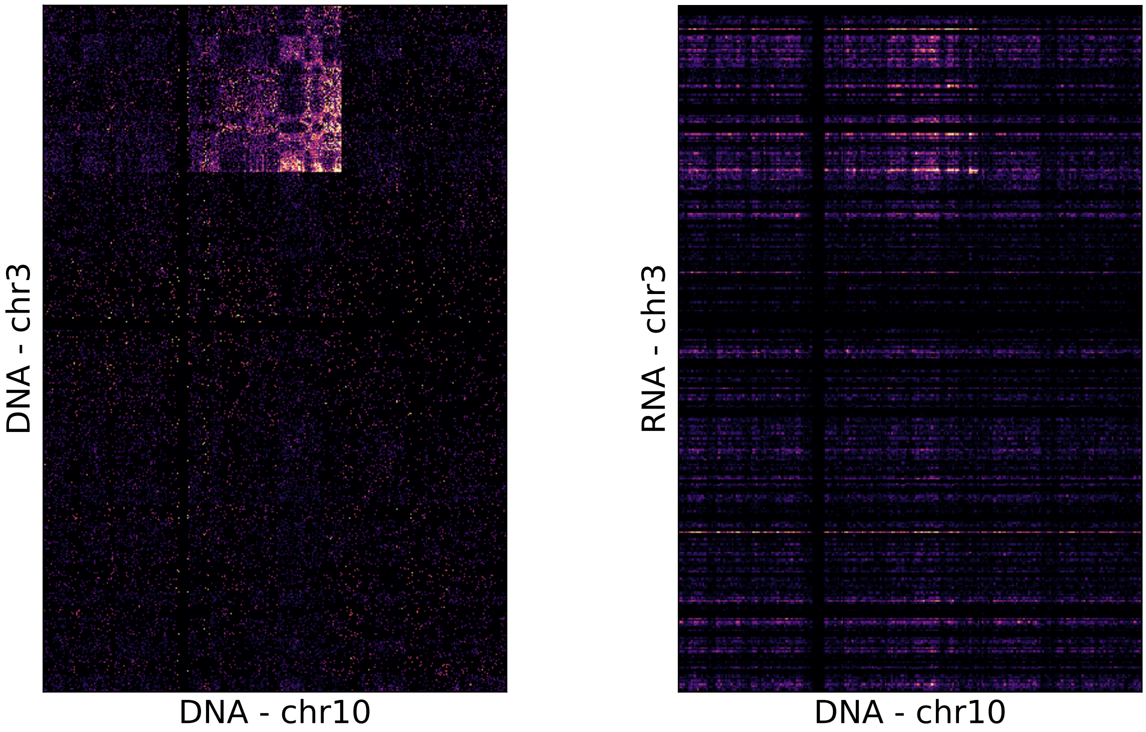

**Figure S7.** An example of the effect of chromosomal rearrangements of the K562 cell line on interactomes: Hi-C data on the left, Red-C data on the right.

**Table S5.** The number of chromosomes (pairs of chromosomes for the interchromosomal case) on which there is an association of contacts with the chromatin structure (statistics were calculated on peaks counts, due to their small width and high coverage of “one-to-all" data).

| RNA | RnaChromDB ID | Number of chromosomes | Number of chromosome pairs |
|---|---|---|---|
| Malat1 | 36 | 18 | 186 |
| Malat1 | 52 | 20 | 178 |
| Malat1 | 94 | 20 | 190 |
| Firre | 198 | 20 | 190 |
| Xist | 156 | 20 | 184 |

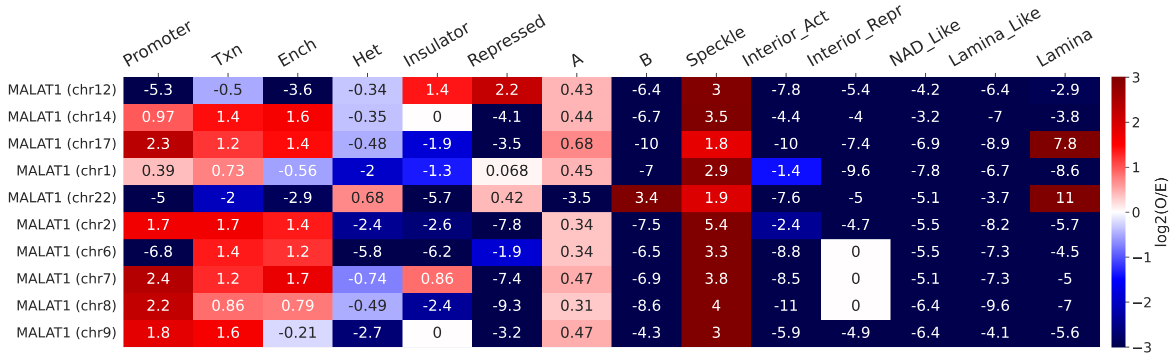

**Figure S8.** The enrichment of paired contacts of MALAT1 RNA in chromatin annotations on different chromosomes.

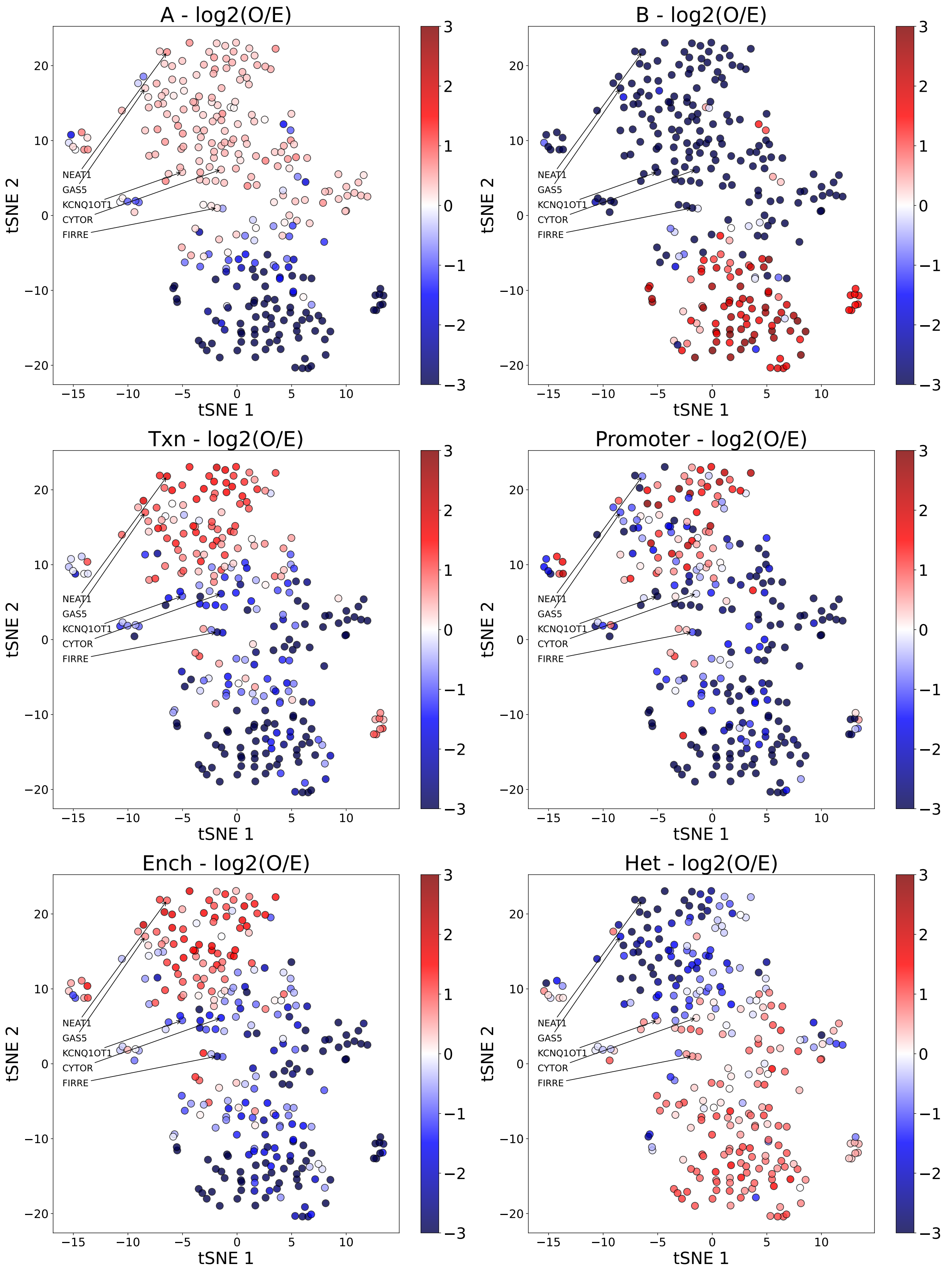

**Figure S9.** tSNE is colored by enrichment of paired contacts of non-coding RNAs in various chromatin state (continued on the next page).

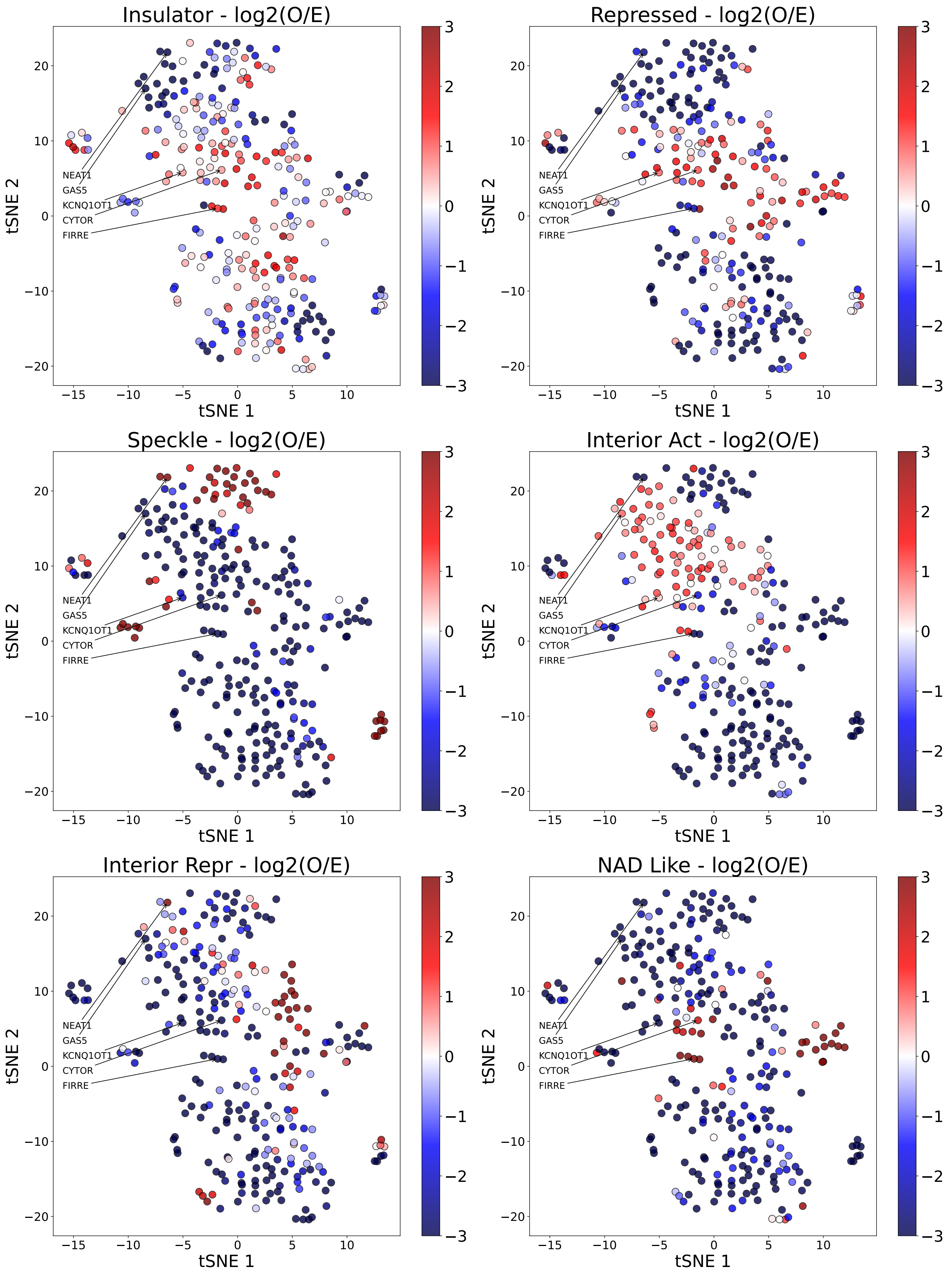

**Figure S9.** tSNE is colored by enrichment of paired contacts of non-coding RNAs in various chromatin state (continued on the next page).

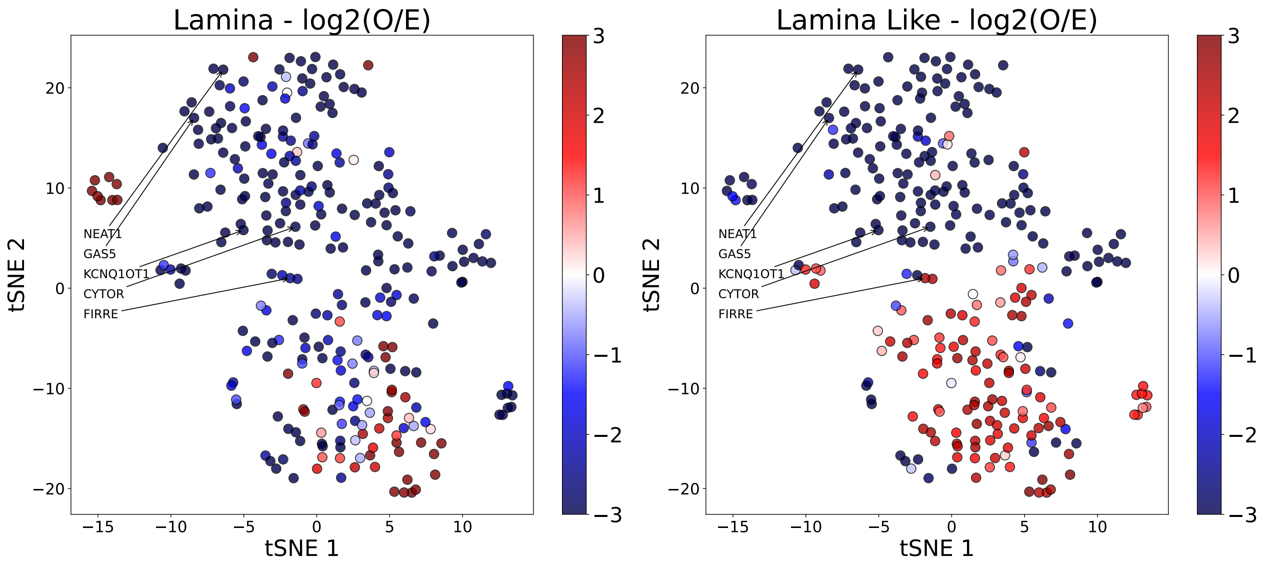

**Figure S9.** tSNE is colored by enrichment of paired contacts of non-coding RNAs in various chromatin state

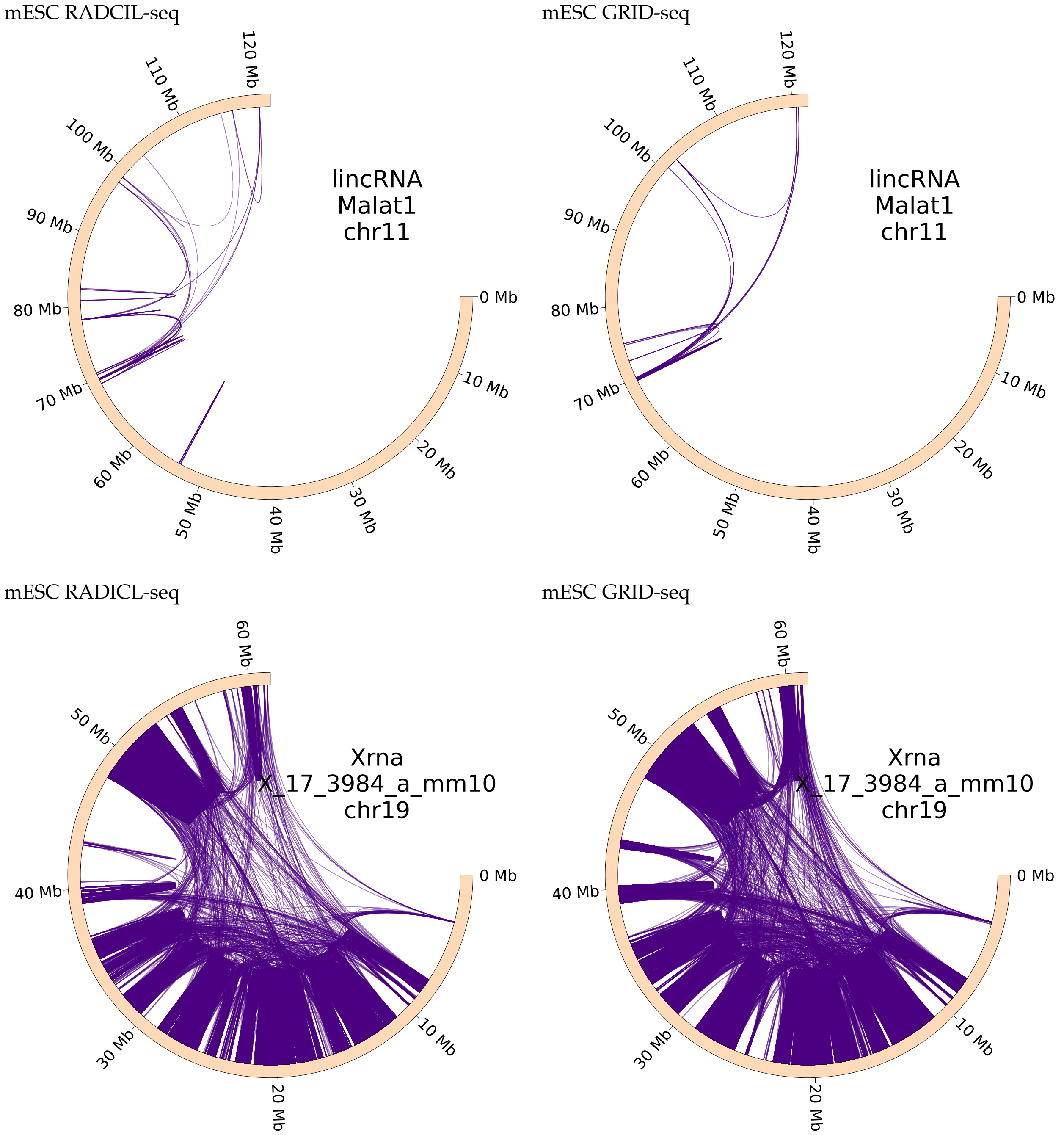

**Figure S10.** Examples of paired contacts of RNA from the protocols RADICL-seq (left) and GRID-set (right) of the mESC cell line. The top row is Malat1. The bottom row is X_17_3984_a_mm10.

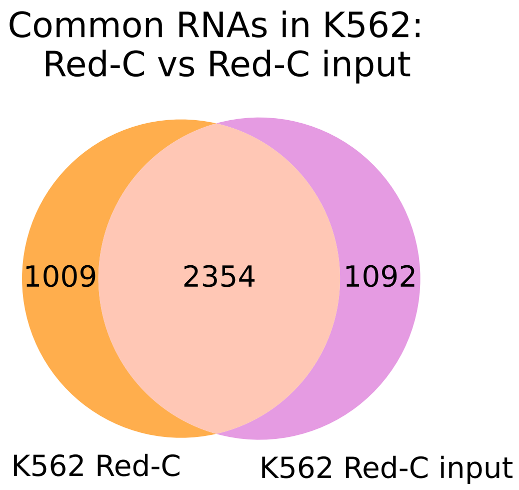

**Figure S11.** The intersection of sets of RNAs associated with the chromatin structure between the Red-C and Red-C (input) protocols, cell line K562, on the left - all RNAs, on the right - non-coding RNAs.

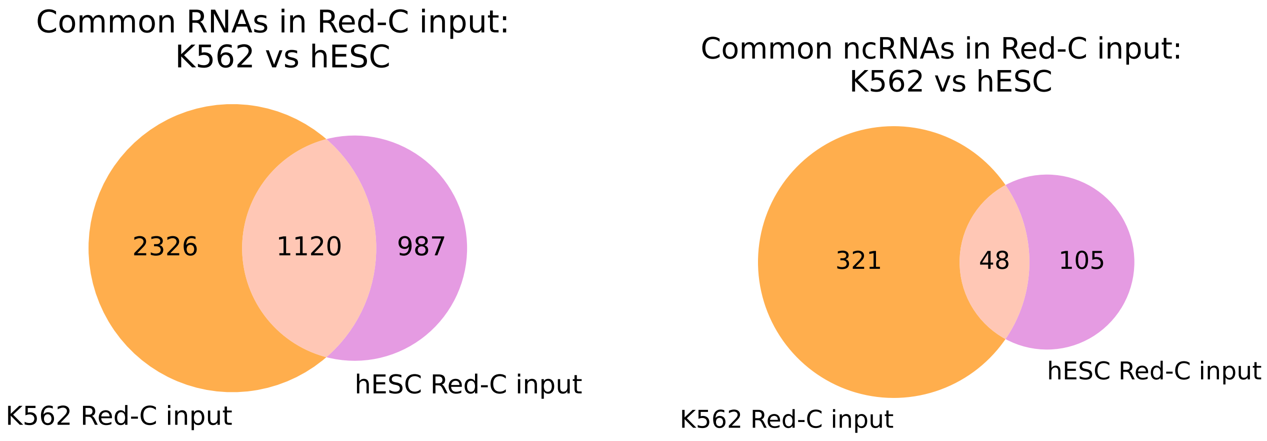

**Figure S12.** The intersection of the sets of RNAs associated with the chromatin structure between the protocols of the K562 and hESC cell lines, the Red-C (input) protocol, on the left - all RNAs, on the right - non-coding RNAs.

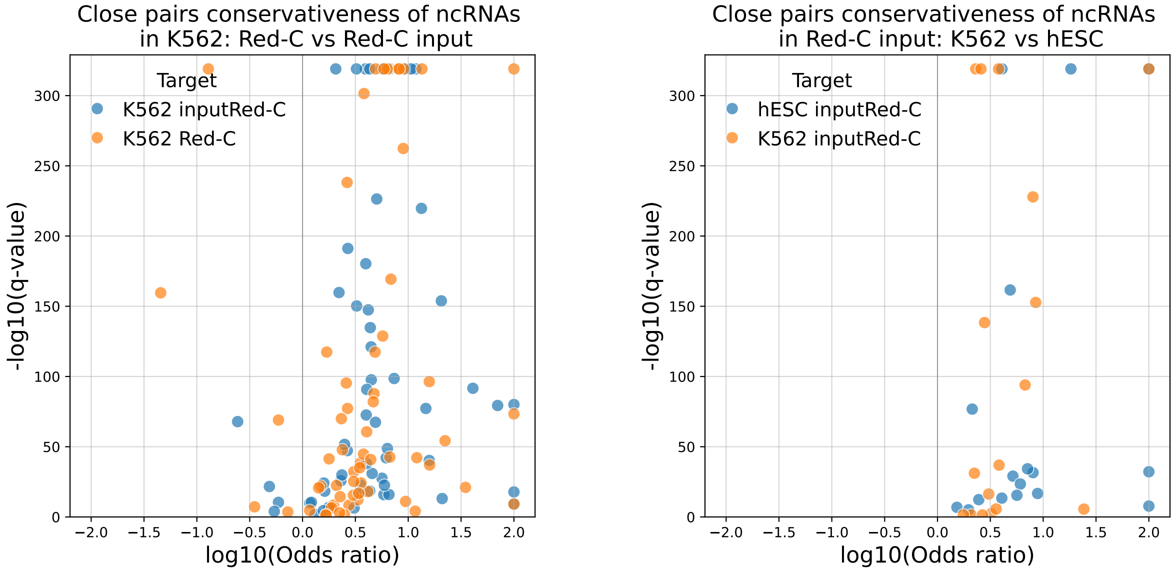

**Figure S13.** The relationship of the q-values and the odds ratios of the Fisher's exact test. On the left - comparison of Red-C and Red-C (input) experiments, on the right comparison of K562 and hESC cell lines.

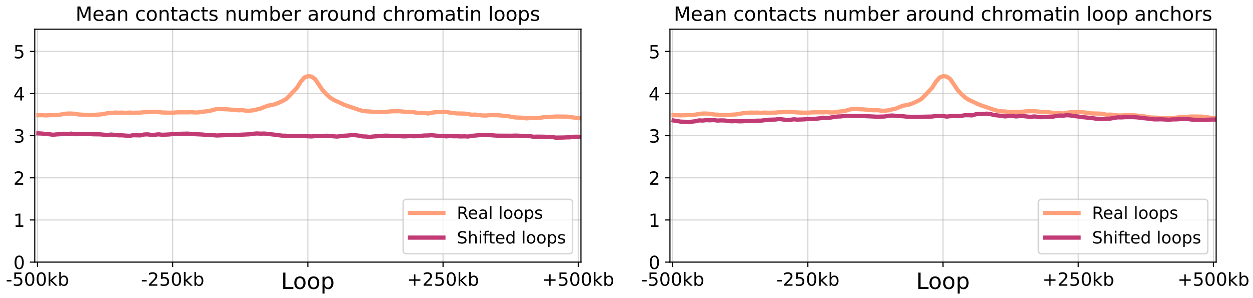

**Figure S14.** The average number of RNA-DNA (K562 Red-C) interactions in the vicinity of chromatin loop anchors, on the left the background model is random shift, on the right the background model is shift, taking into account belonging to the A/B compartment.

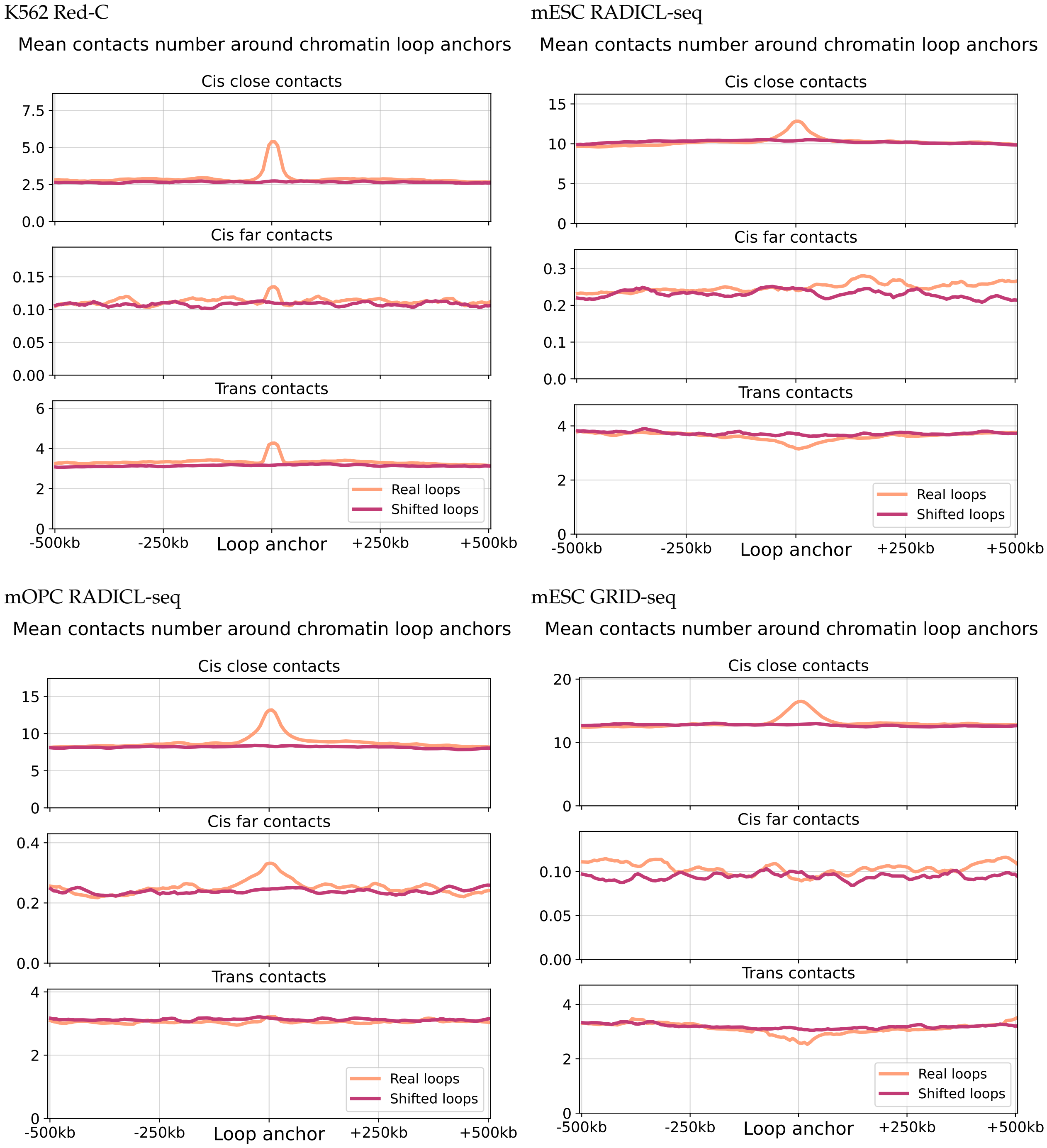

**Figure S15.** The average number of RNA-DNA interactions in the vicinity of chromatin loop anchors (continued on the next page).

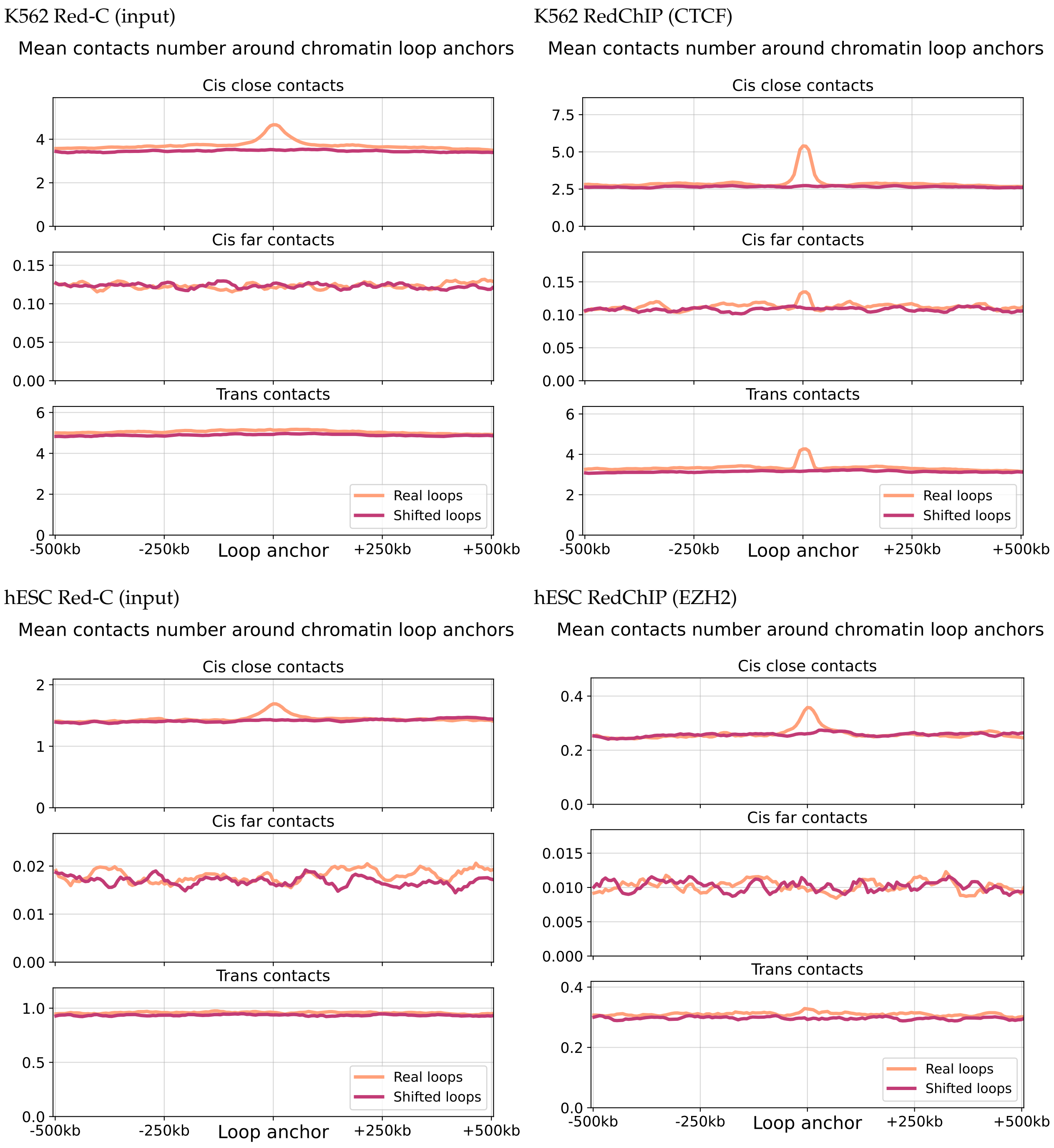

**Figure S15.** The average number of RNA-DNA interactions in the vicinity of chromatin loops anchors.

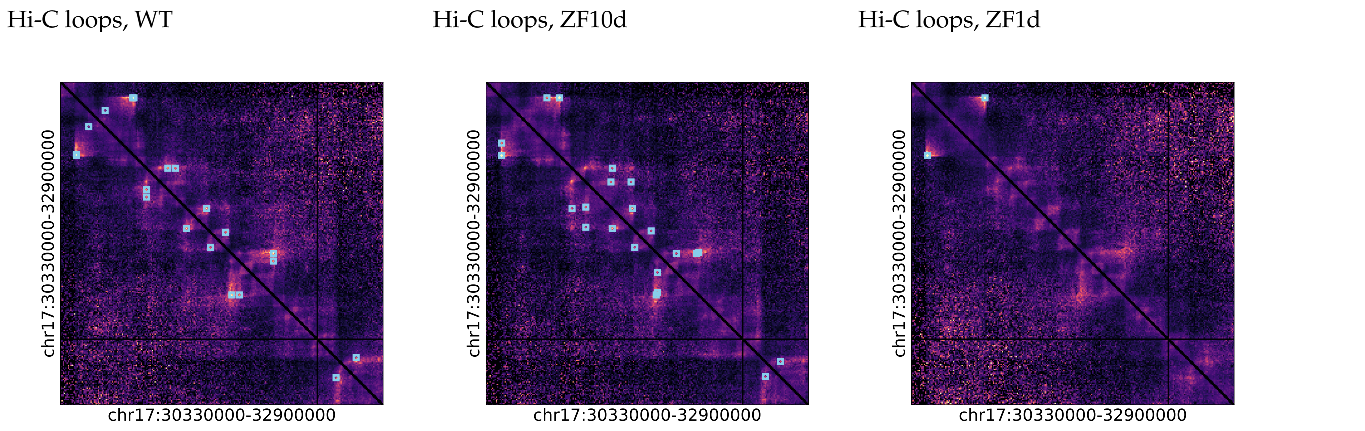

**Figure S16.** Examples of annotated loops on Hi-C maps in different conditions, on the left - WT, in the center - deletion in ZF10, on the right - deletion in ZF1 (chr17:30330000:32900000).

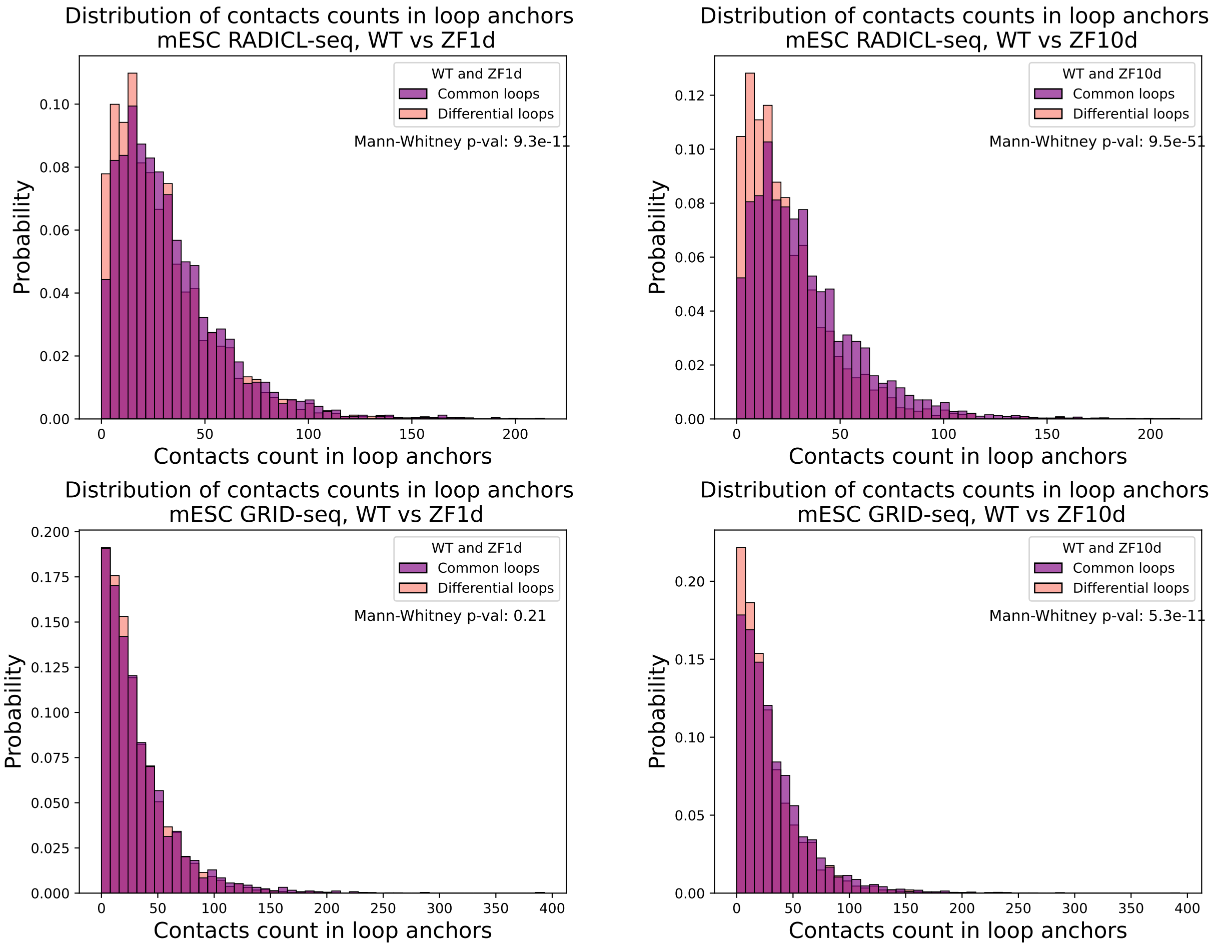

**Figure S17.** Comparison of the distribution of the number of RD-contacts in mESC within the anchors of chromatin loops. Common loops are found in both WT and mutant lines. Differential loops are absent in the mutant cell line (on the left is a deletion in the ZF1 domain, on the right is in the ZF10 domain, the upper row is RADICL-seq, the lower one is GRID-seq).

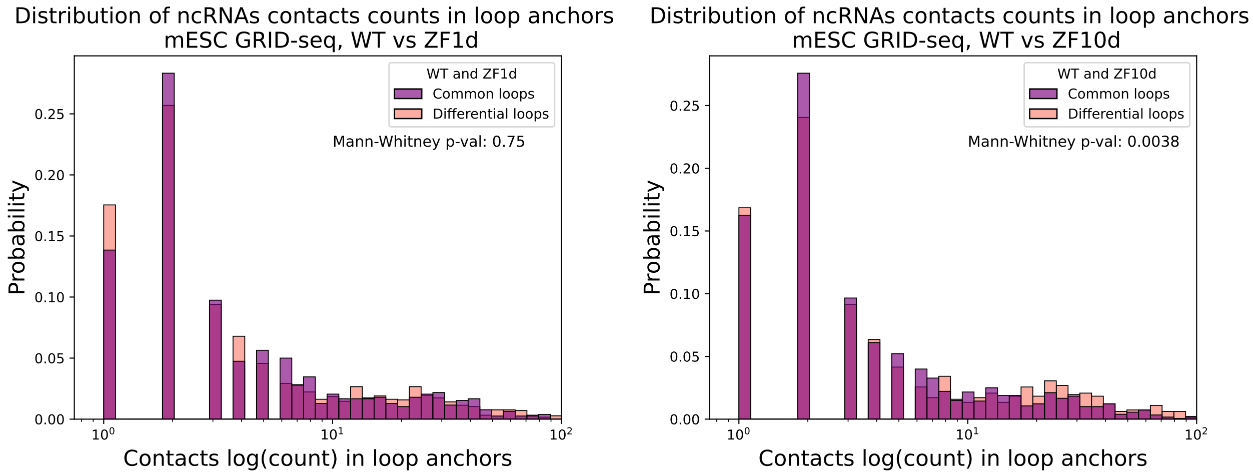

**Figure S18.** Comparison of the distribution of the number of ncRNAs RD-contacts in mESC GRID-seq within the anchors of chromatin loops. Common loops are found in both WT and mutant lines. Differential loops are absent in the mutant cell line (on the left is a deletion in the ZF1 domain, on the right is in the ZF10 domain.

mESC RADICL-seq

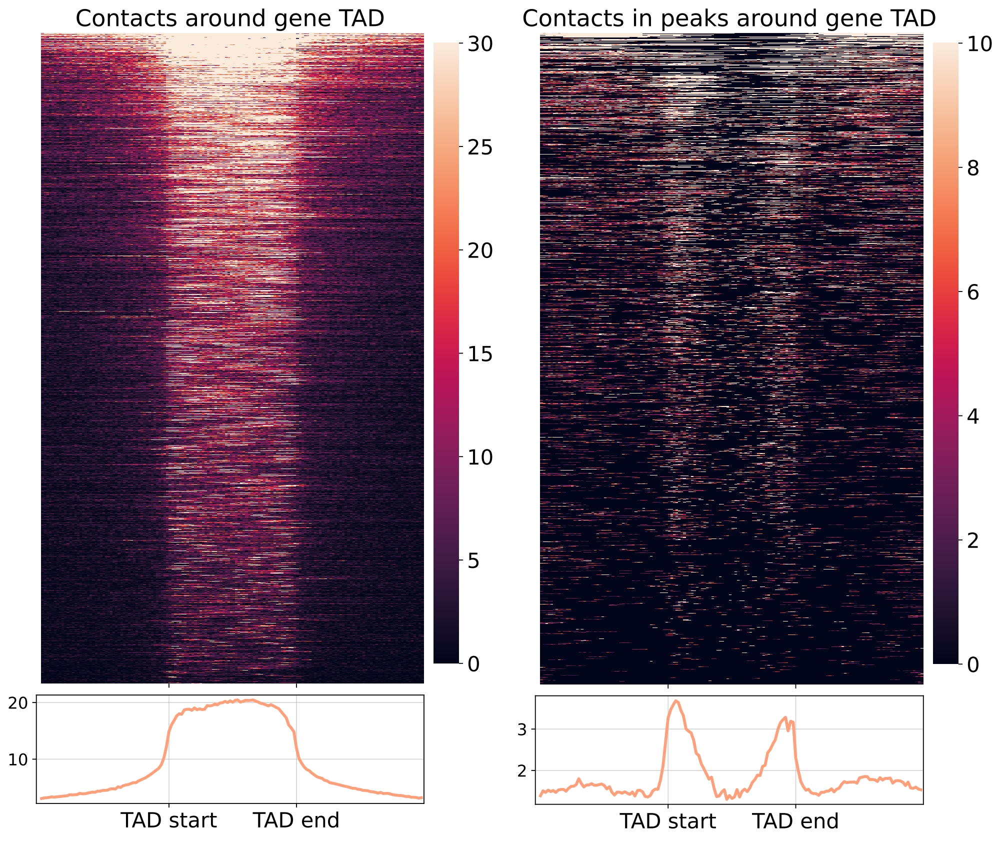

mOPC RADICL-seq

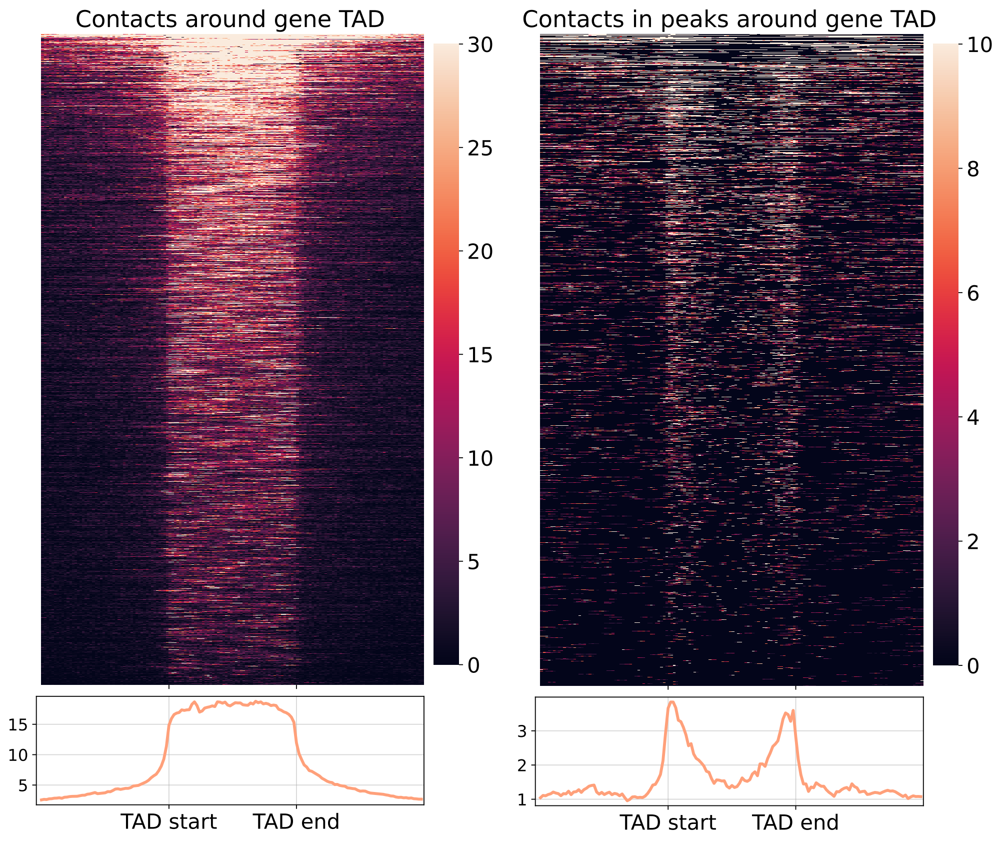

**Figure S19.** Left - distribution of the full set of RD-contacts around the TADs of RNAs genes (Red-C K562), right - distribution of RD-contacts from BaRDIC peaks around the TADs of RNAs genes (continued on the next page).

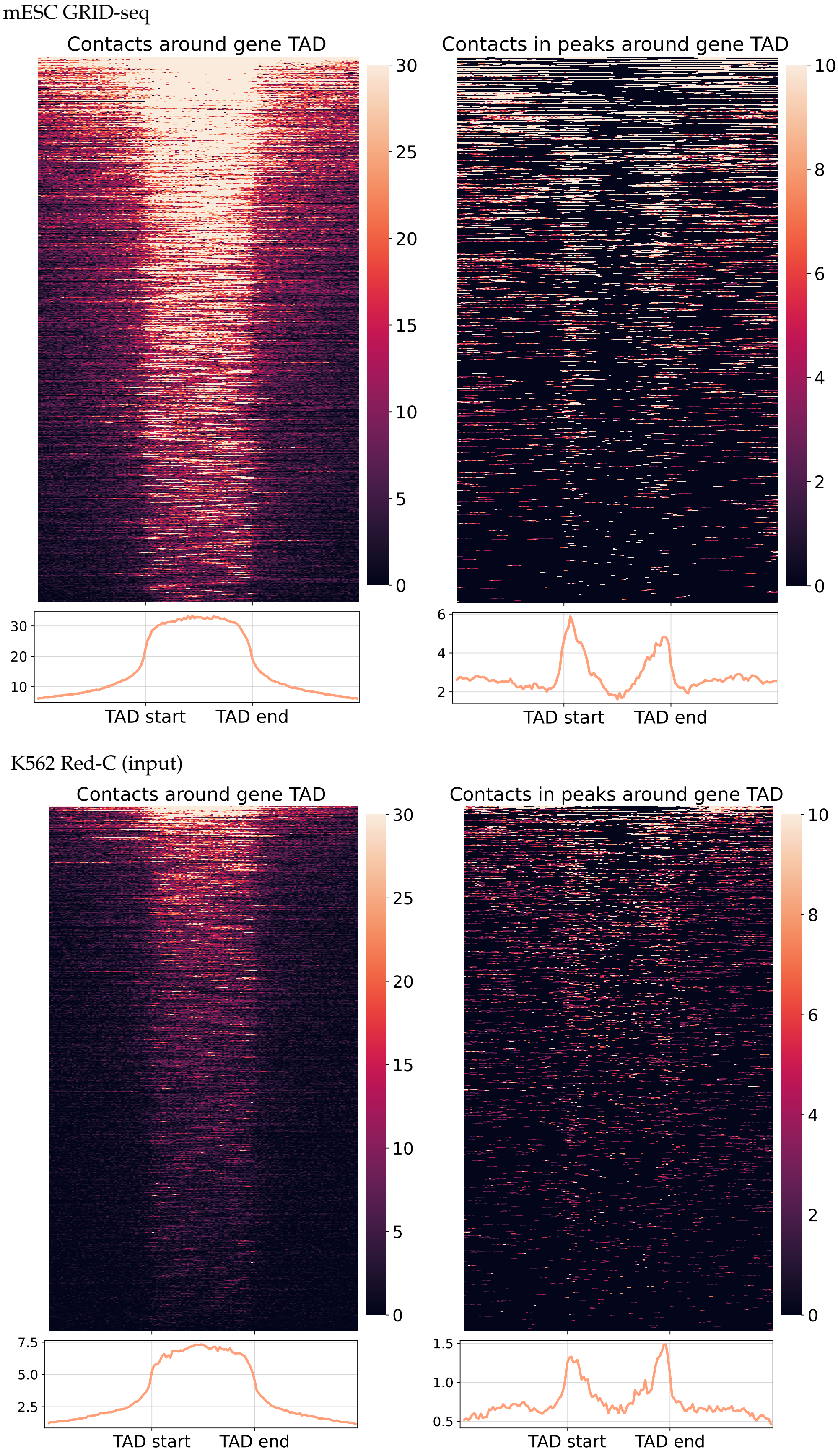

**Figure S19.** Left - distribution of the full set of RD-contacts around the TADs of RNAs genes (Red-C K562), right - distribution of RD-contacts from BaRDIC peaks around the TADs of RNAs genes (continued on the next page).

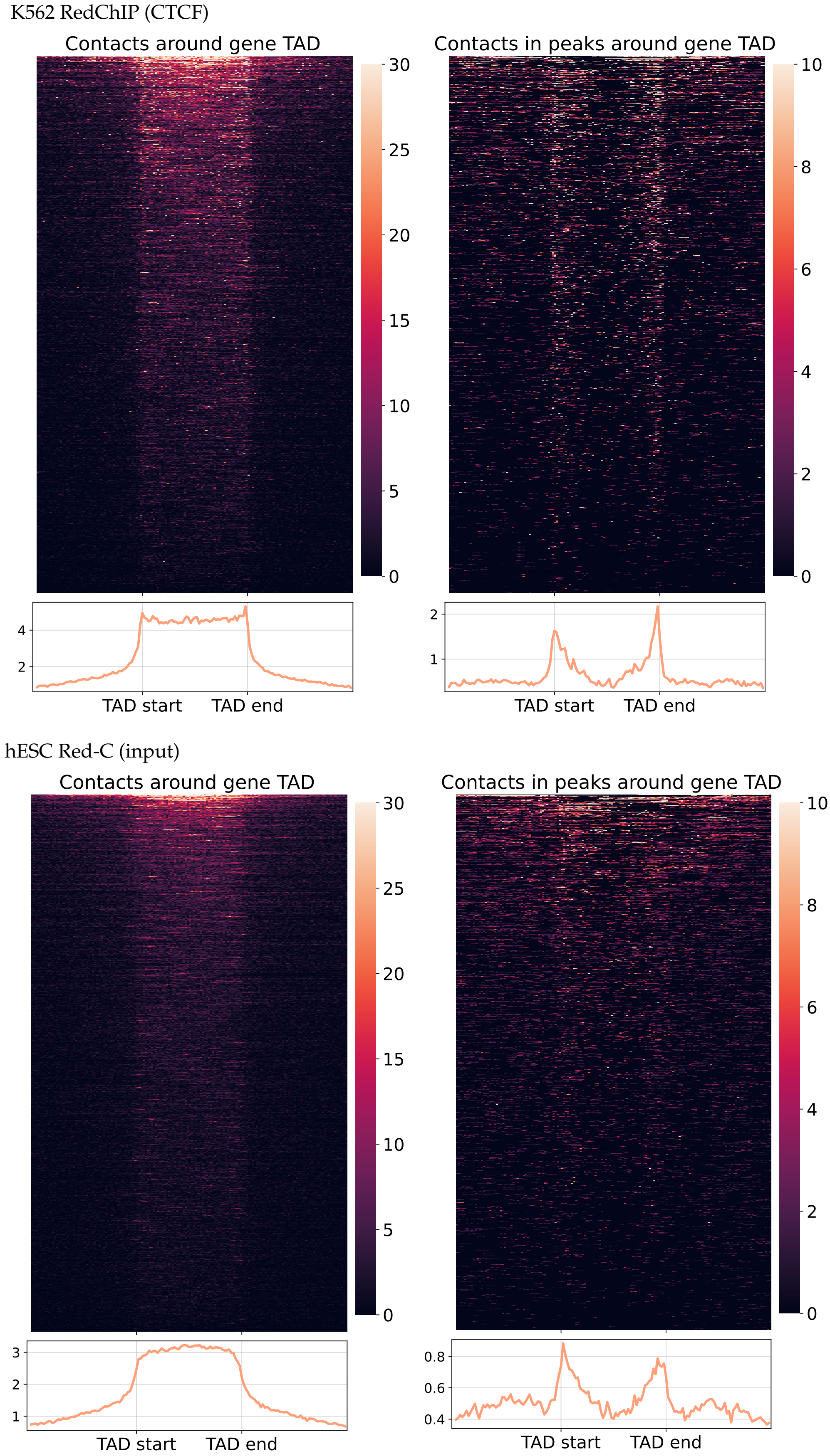

**Figure S19.** Left - distribution of the full set of RD-contacts around the TADs of RNAs genes (Red-C K562), right - distribution of RD-contacts from BaRDIC peaks around the TADs of RNAs genes (continued on the next page).

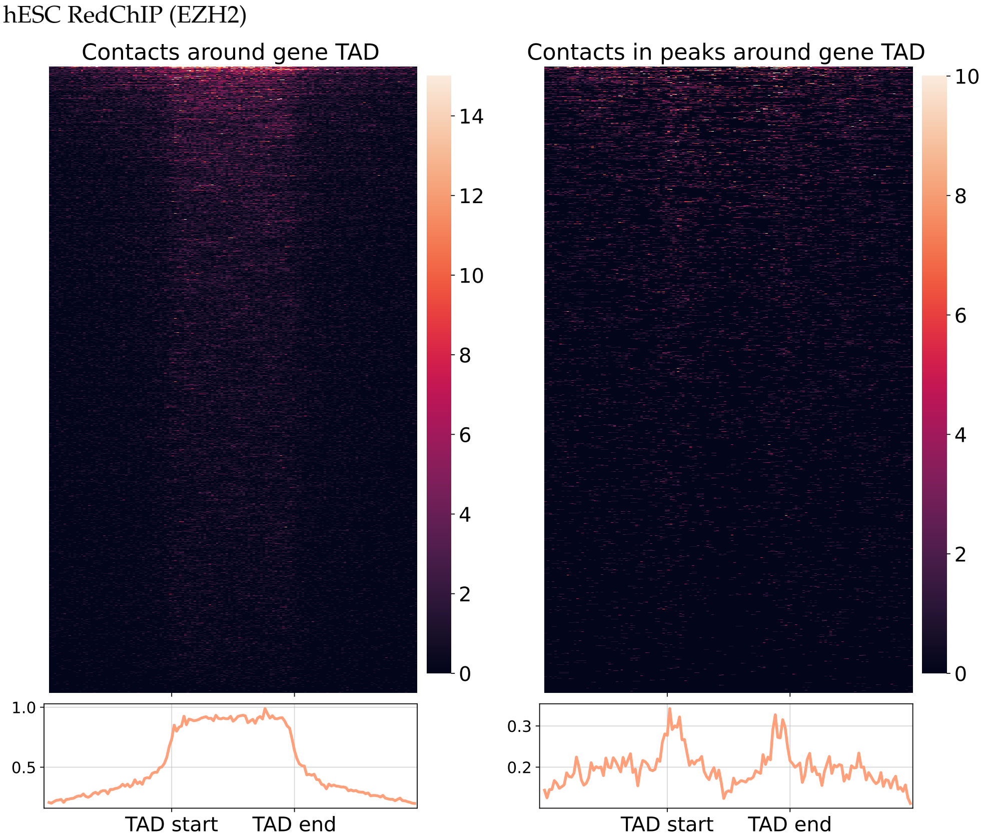

**Figure S19.** Left - distribution of the full set of RD-contacts around the TADs of RNAs genes (Red-C K562), right - distribution of RD-contacts from BaRDIC peaks around the TADs of RNAs genes.

**Table S6.** A two-sided Mann-Whitney test for OutInDR values of different RNA biotypes and mRNA (NS - non significant, adj p-value > 0.05).

| Cell line | RNA type | Mann-Whitney p-value (Bonferroni correction) |
|---|---|---|
| K562 Red-C | lncRNA | 3.7e-08 |
| | vlincRNA | 2.23e-21 |
| | pseudogene | NS |
| | xRNA | NS |
| K562 Red-C (input) | lncRNA | 9.90e-07 |
| | vlincRNA | 8.46e-11 |
| | pseudogene | NS |
| | xRNA | NS |
| hESC Red-C (input) | lncRNA | NS |
| | vlincRNA | NS |
| | pseudogene | NS |
| | xRNA | NS |
| mESC RADICL-seq | lincRNA | NS |
| | antisense | NS |
| | processed transcript | NS |
| | xRNA | 4.24e-09 |
| | TEC | NS |
| mOPC RADICL-seq | lincRNA | NS |
| | antisense | NS |
| | processed transcript | NS |
| | xRNA | 2.60e-13 |
| | TEC | NS |
| mESC GRID-seq | lincRNA | 0.032 |
| | antisense | NS |
| | processed transcript | NS |
| | xRNA | 7.33e-11 |
| | TEC | 2.60e-08 |

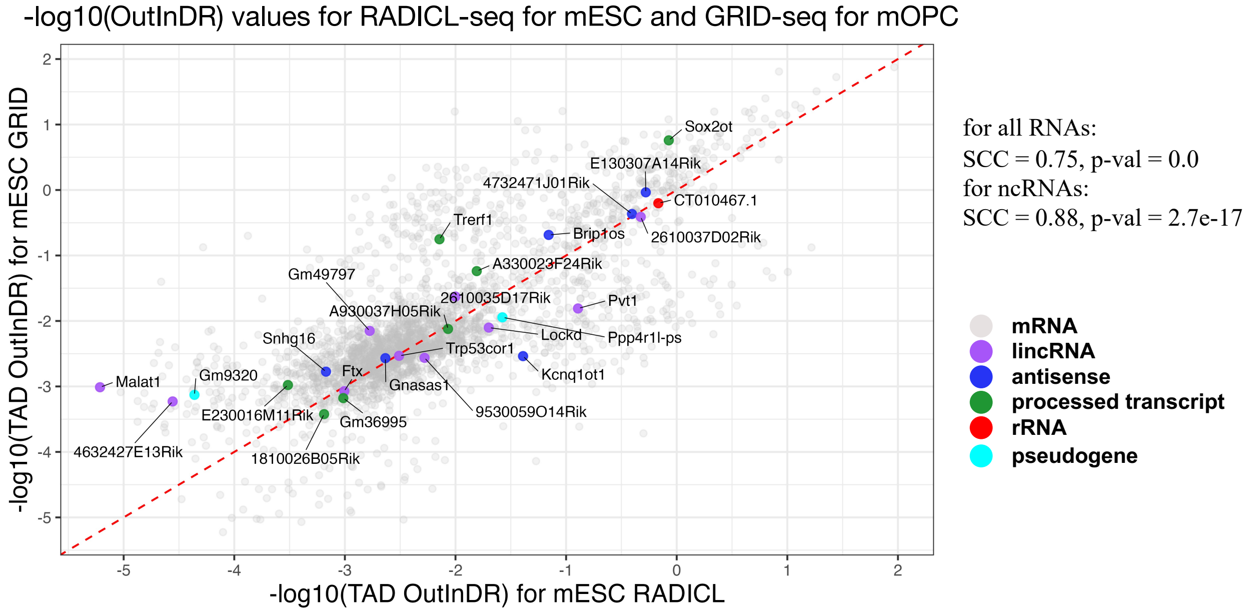

**Figure S20.** Scatterplot with values -log10(OutInDR) for the RADICL-seq experiment on the X axis and GRID-seq on the Y axis for the mESC cell line.

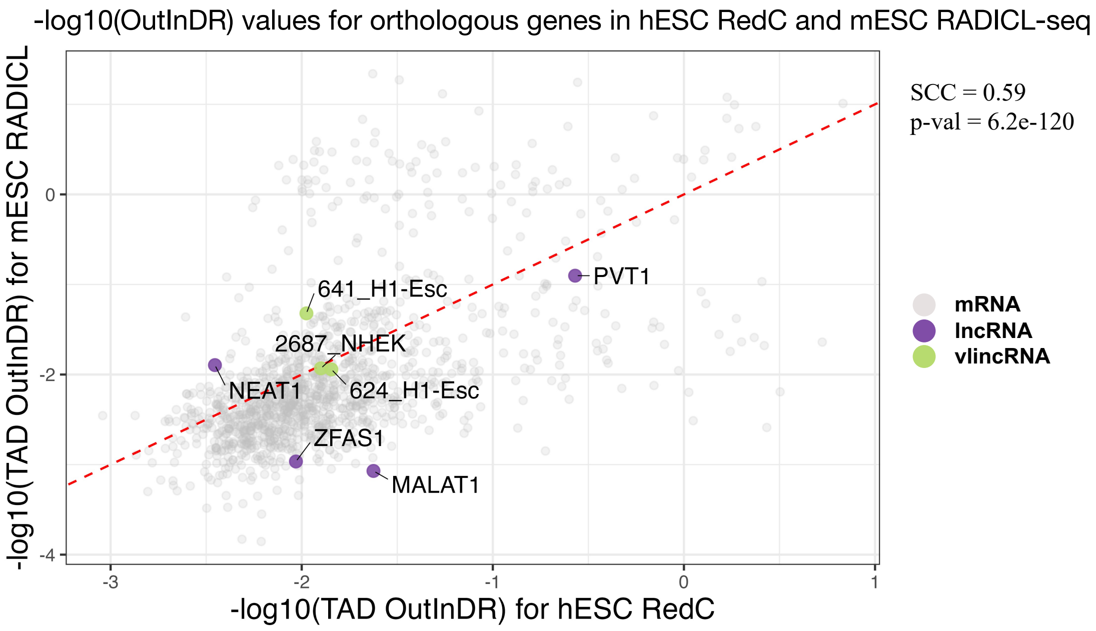

**Figure S21.** Scatterplot with values -log10(OutInDR) of orthologous genes in the hESC (X-axis) and mESC RADICL-seq (Y-axis) cell lines.

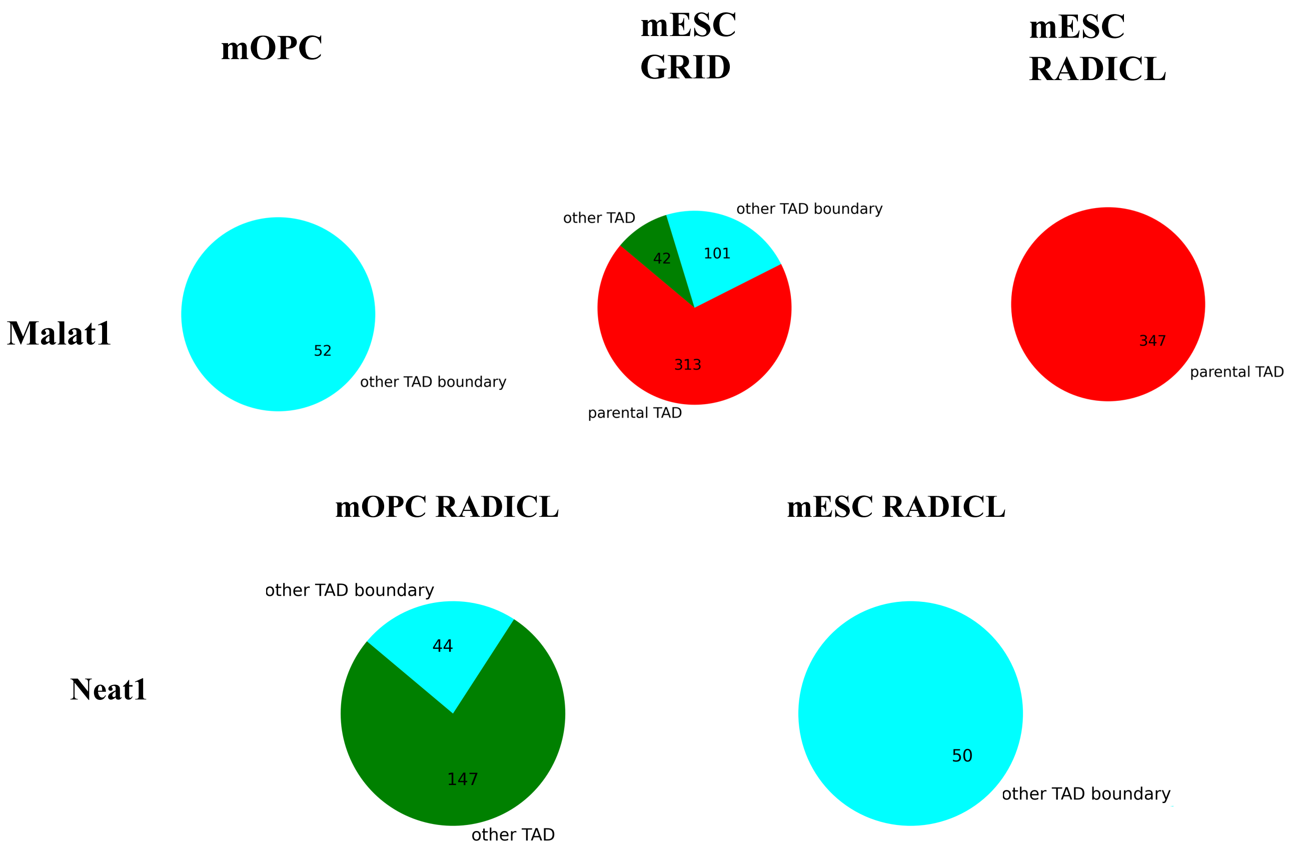

**Figure S22.** RD-contacts distribution in TADs and boundaries in different cell lines. Top row - MALAT1 contacts, bottom row - NEAT1.

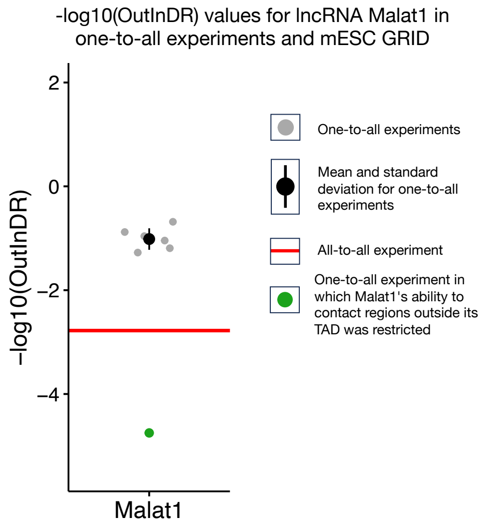

**Figure S23.** Comparison of the -log10(OutInDR) value for lincRNA Malat1 in "one-to-all" and "all-to-all" experiments for mESC GRID-seq.

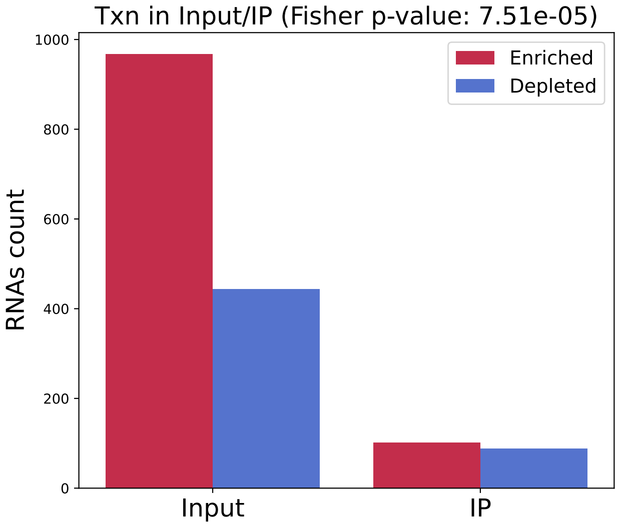

**Figure S24.** The number of cases of significant overrepresentation and underrepresentation of paired contacts in pairs of the active transcription state in Red-C and RedChIP.

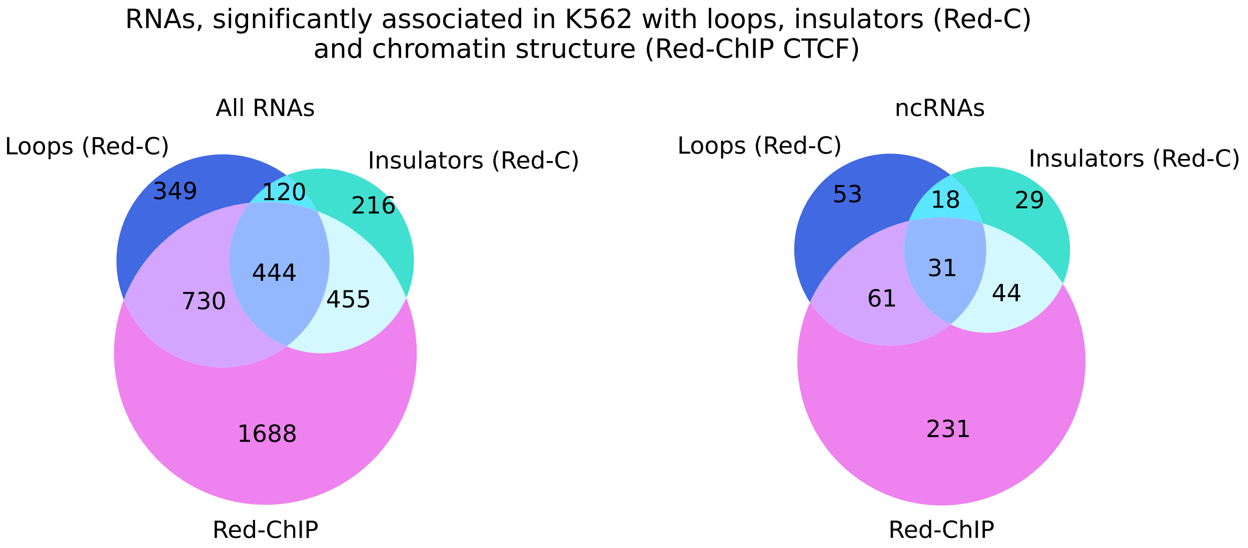

**Figure S25.** The intersection of RNAs which are significantly associated with loops (Red-C data), with the chromatin structure (Red-ChIP), and whose paired contacts are associated with insulators (Red-C).
